# Colocalization features for classification of tumors using desorption electrospray ionization mass spectrometry imaging

**DOI:** 10.1101/440057

**Authors:** Paolo Inglese, Gonçalo Correia, Pamela Pruski, Robert C Glen, Zoltan Takats

## Abstract

Supervised modeling of mass spectrometry imaging (MSI) data is a crucial component for the detection of the distinct molecular characteristics of cancerous tissues. Currently, two types of supervised analyses are mainly used on MSI data: pixel-wise segmentation of sample images, and whole-sample-based classification. A large number of mass spectra associated with each MSI sample can represent a challenge for designing models that simultaneously preserve the overall molecular content while capturing valuable information contained in the MSI data. Furthermore, intensity-related batch effects can introduce biases in the statistical models.

Here we introduce a method based on ion colocalization features that allows the classification of whole tissue specimens using MSI data, which naturally preserves the spatial information associated the with the mass spectra and is less sensitive to possible batch effects. Finally, we propose data visualization strategies for the inspection of the derived networks, which can be used to assess whether the correlation differences are related to co-expression/suppression or disjoint spatial localization patterns and can suggest hypotheses based on the underlying mechanisms associated with the different classes of analyzed samples.

## INTRODUCTION

Mass spectrometric imaging (MSI) has been widely applied to investigate the spatial distributions of molecular species within tissue specimens. In particular, it has shown promising results to serve as a technological platform for the study of chemical heterogeneity of samples. Particularly in the case of cancer tissue, established protocols exist that are tailored to the imaging analysis of human biopsy specimens (either freshly prepared - frozen tissue - or - formalin fixed paraffin embedded (FFPE), etc. - ^1,2^) using either desorption electrospray ionization (DESI) ^3,4^, matrix-assisted laser desorption ionization mass spectrometry (MALDI-MS) ^5–10^ or secondary ionization mass spectrometry (SIMS) ^11–14^ as the predominant MSI analytical techniques. Applying any of these techniques to a sample generates a set of ion images, which we denote as mass spectrometry (MS) image, or equivalently, MSI sample in this text.

Two major statistical modelling approaches are often employed to identify relationships between ion signatures and sample properties of interest. Unsupervised methods try to capture the intrinsic statistical properties of the MSI data, such as spectral similarity and generate partitions of the data without relying on any external ground truth. These methods can be useful to suggest novel hypotheses on the metabolic pathways associated with cancerous tissue, for instance, when histological or other expert-driven properties of the analysed samples are missing^15–20^. In contrast, supervised methods aim to determine statistical relationships between the observed ion signatures and ‘labels’ associated with the analysed data, that are manually generated by experts. Two different types of supervised modelling can be applied to MSI data. The first approach, called ‘segmentation’ uses the manual labels associated with pixel spectra to partition the MS images into regions characterised by the same property of interest^21^. For instance, a segmentation method can predict the labels of MS image pixels, generating a spatial map of regions associated with tumor, necrotic, or surrounding connective tissue^22–24^. The second approach aims to predict the label of an entire MS image, based on the molecular information contained in it. For example, a model can be trained to discriminate between MS images from healthy tissue sections and cancerous tissue sections, or between different types of cancer.

The importance of these types of models is evident considering that they allow associating a general property to the entire MSI sample, with the possibility of determining general molecular patterns that are responsible for the different classes of analysed samples.

However, the large number of mass spectra present in a single MSI sample makes it challenging to design a model that captures a single property (the ‘label’) preserving its molecular information. Currently, the most used approach is based on extracting a ‘representative’ spectrum for the entire MS image, which is then related to the corresponding label^25,26^. A typical choice is the mean spectrum calculated over pixels of the same tissue type. Another approach is based on selecting a random subset of pixel spectra from the tissue of interest and predicting their class^27^.

Clearly, these approaches have limitations. The first approach fails to preserve the two most important properties of MS images, their molecular heterogeneity and spatial distributions, compressing all the data into a single spectrum. The second approach introduces new challenges in determining the final prediction of the entire MS image. For instance, if the selected pixels are predicted to belong to different classes, how is the final prediction (that is related to the entire MS image) generated?^18^.

Another difficulty associated with all these methods is that they require the precise annotation of a histologically homogeneous region, from where the pixels are extracted and associated with the global sample label.

Ultimately, statistical models can be biased by the presence of batch effects. These may originate from several (often difficult to control) sources, such as instrumental conditions (for instance, temperature fluctuations, drifts in mass/charge ratio calibration), or sample-related properties (for example, tissue chemical matrix, ion suppression)^28^. Biased datasets (consisting of multiple MSI samples) often exhibit systematic variations of the mass spectral intensities of specific MSI samples, that are not a product of their inner properties, but a consequence of the specific experimental conditions during the spectral acquisition.

For this reason, normalization techniques are necessary to reduce the spectral intensity variations due to possible batch effects^29^.

Here we present an alternative approach for the classification of entire MSI samples that addresses the two challenges discussed, preserving more of the rich molecular information and being less sensitive to the systematic variations of mass spectral intensities induced by batch effects.

The hypothesis behind the method presented for classifying tumor MSI images is that the observed spatial patterns of the ion peak intensities reflect the metabolic properties of the analyzed sample. This hypothesis is compatible with the typical observation in the supervised segmentation of MS images, where a set of ions are characterized by a significantly higher (lower) peak intensity in a specific region of the tissue section. This is equivalent to saying that these ions are more (less) colocalized in that area than in the remaining part of the tissue. Additionally, the central assumption underlying this type of analysis is that the expression of these ions relates to either one or more metabolic mechanisms occurring within the local region of the sample.

Here, we apply the concept of ion colocalization to represent the entire molecular information of individual tissue samples. In this way, the whole sample can be expressed in its entirety by a single vector of colocalization features. Additionally, possible issues caused by batch-to-batch variability^30,31^ can be mitigated, since colocalization is a relative measure and, therefore, is less sensitive to systematic variations of ion intensities across multiple samples. A similar concept has been previously employed for unsupervised (in contrast to supervised in this work) analysis of MSI data (generating clusters of highly colocalized ions)^32–36^.

Conceptually, the MSI data is converted into a graph, and then network features are used for supervised classification, analogously to Amoroso et al.^37^, where the complex networks framework is employed for classifying patients with neurodegenerative disease. Specifically, Pearson’s correlation has been used as a suitable measure for determining the degree of colocalization, in particular in fluorescence microscopy^38^.

In this paper, we show how the colocalization features can be used to classify three types of cancer specimens (breast, ovarian and colorectal). This approach results in higher predictive performance potentially providing a deeper insight into the possible mechanisms underlying their biochemical differences than the current state-of-the-art methodologies.

## MATERIALS AND METHODS

Sixty-nine specimens from three distinct cohorts of human tumors were cryo-sectioned and subsequently analyzed by DESI-MSI. The full dataset consisted of twenty-eight breast cancer, part of the dataset published in Guenther et al.^26^, sixteen colorectal adenocarcinomas, part of the dataset published in Inglese et al.^36^ and twenty-five ovarian cancer samples, part of the dataset published in Doria et al.^39^. All the raw spectra were acquired in the negative ion mode using a custom DESI sprayer ion source interfaced to a high-resolution orbital trapping mass spectrometer [Exactive, Thermo Fisher Scientific] in the range of 150 – 1,000 *m/z* using the experimental and instrumental conditions reported in the Supplementary Table S1. For the data analysis, the tissue sections were split into two random subsets, denoted *cross-validation set* and *test set* consisting of sixteen tissue sections per tumor type, and twelve breast plus nine ovarian cancer sections, respectively.

### Pre-processing

All the raw spectral data were first converted into imzML format (centroided mode)^40^ and their *m/z* values were corrected using a single point re-calibration, using a locally estimated scatterplot smoothing (LOESS) model fitted on the distance between the observed and the theoretical *m/z* values of palmitic acid (255.2330 *m/z*, [M-H]^−^)^36^. Observed *m/z* values corresponded to the closest peak within a search window of 10 ppm, compatible with the theoretical instrumental error. The use of LOESS allowed correcting for the global *m/z* drift trends while preserving the small fluctuations.

The re-calibrated image samples were independently pre-processed through a workflow consisting of a) total-ion-count (TIC) scaling intensity normalization, b) peak matching, c) log-transformation using MALDIquant for R^41,42^. The choice of TIC-scaling normalization was based on the fact that we were interested in the colocalization patterns based on the relative abundance of the spectral peaks. TIC-scaling also simplified the interpretation of the results, where the normalized ion intensities could be read as fractions of the total amount of detected molecules in each pixel. The peak matching procedure assigned the detected peaks intensity from each pixel of an MS image to a *common m/z* vector. Each pixel spectrum was allowed to contribute to a specific *common m/z* value with only one peak (Supplementary Table S2). No inter-sample normalization to reduce possible inter-samples batch effect was applied.

Tissue-unrelated peaks were filtered using the global reference and pixel count filters of the SPUTNIK package for R^43^. The parameters used are reported in Supplementary Table S3.

In order to match the common peaks detected in the MS images, a similar peak matching procedure was applied to the spectra representative of the MS images. The average of the non-zero peaks intensity within each MS image was used as a representative spectrum of each MS image of the *cross-validation set*. Only the matched peaks detected in all the images were retained for the statistical modeling (Supplementary Table S4). In this way, we ensured that the different colocalization patterns among the tumor types involved the molecules observed in all the samples. This strict constraint was necessary because ion presence/absence patterns may be a consequence of non-biologically-related mechanisms (e.g. different tissue chemical matrices).

Isotopes were removed from the *common m/z* vector. These were identified as common *m/z* values that fell in the interval [*m* − *k × 1.002, m + k × 1.0045*], where 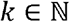 and *m* is a member of the common *m/z* vector^36^. Manual inspection was performed to identify and remove possible isotopes left after the de-isotoping procedure. The de-isotoping procedure aimed to reduce the number of the final features, removing those characterized by a redundant molecular information (isotopes are expected to equally colocalize with their monoisotopic forms).

The *test set* peaks were matched with the common *m/z* vector of the *cross-validation set* within a search window of ±5 ppm. When multiple peaks matched, the *m/z* value of the most intense peak was selected. If no matched peak was found, the intensity was left equal to 0 in all the pixels. This procedure was intended to evaluate the performances of the classification model with out-of-sample MS images (i.e. MS images that are not part of the dataset used for training the models).

### Colocalization features extraction

The MSI images comprised both tissue-related and tissue-unrelated (off-tissue areas) pixels. In order to use the only tissue-related pixels for the calculation of the colocalization features, a binary mask (region of interest, ROI) representing the area of the image occupied by the tissue was determined by applying the ‘*kmeans2*’ method from the SPUTNIK package for R^43^. Four clusters were extracted by k-means applied to the entire MSI images, with the purpose of identifying a finer segmentation of the MS image onto tissue-related and tissue-unrelated areas. In this way, the tissue-related spectra characterized by signal intensity close to the off-tissue areas could also be distinguished, providing a more precise localization of the area occupied by the tissue. The clusters that were not localized in the corners of the image were merged to define the tissue-related ROI. This heuristic rule was determined after visual inspection of the segmented images (Supplementary Figure S1).

Spearman’s rank correlations between each pair of matched peaks were calculated within each MS image using the only tissue-related ROI pixels (Supplementary Equations 4-5). Being a rank-based quantity, Spearman’s rank correlation is more robust to the presence of outliers, and it is insensitive to the absolute values of the variables, which are also equivalent to systematic variations.

The asymptotic *t* approximation^44^ was applied to determine the significance of each correlation. The correlations associated with a Benjamini-Hochberg corrected p-value larger than 0.05 were set equal to 0. Subsequently, the elements of the upper triangular correlation matrix (corresponding to the correlations between different pairs of spectral peaks) were vectorized and used as features for the image sample (Figure 1A).

**Figure 1.**
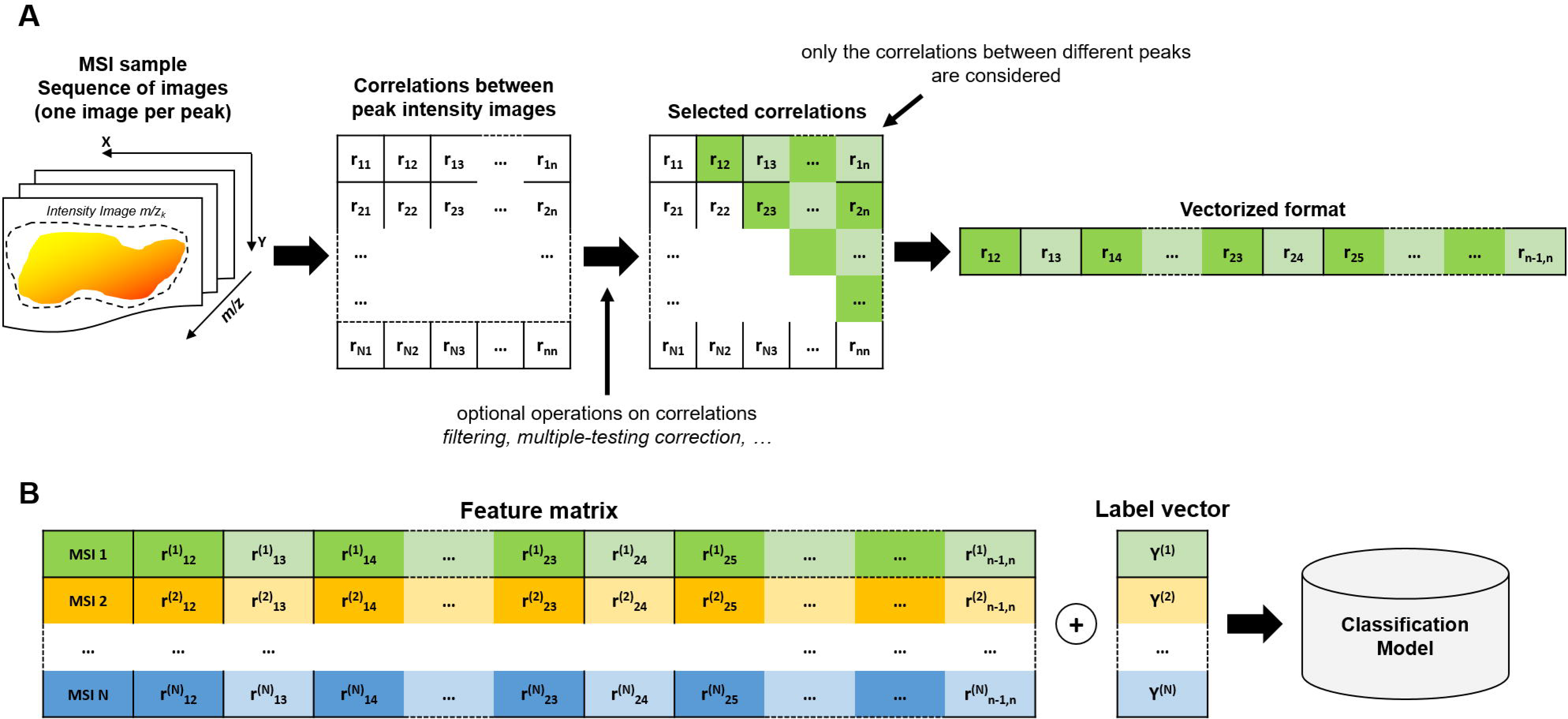
Colocalization based classification scheme. (A) The correlation matrix is calculated from all the pairs of the MS ion images. The upper triangular matrix elements are vectorized to define the feature vector representing the MSI sample. (B) A classification model can be fitted on the colocalization feature vectors and the corresponding MSI sample labels.

Since the size of the tissue-related ROI could differ from one MS image to another, the Spearman’s rank correlations were calculated on a fixed number of randomly selected pixels belonging to the tissue-related ROI. Various sample sizes *N*_*pix*_ = {100, 200, 300, …, 1000} were tested.

No smoothing or other spatially-related intensity transformations were applied to the ion images before calculating the correlations.

In the entire text, the terms colocalization and correlation are treated as synonyms.

### Supervised modeling

The purpose of the supervised model was to predict the tumor type from the colocalization features of an off-sample mass spectrometry image (Supplementary Methods 2). Partial Least Squares – Discriminant Analysis (PLS-DA) algorithm was employed to fit the supervised models on the colocalization features.

Given a fixed number of randomly selected pixels *N*_*pix*_ used for the extraction of the colocalization features, the performance of the classifiers was evaluated through a 10-fold cross-validation scheme applied to the *cross-validation set* of MS images. In each round of the cross-validation, the MS images belonging to the *cross-validation set* were split into a *training* and *validation sets*, and a PLS-DA model was fitted on the ordered pairs of vectors (*f*_*i*_, *Y*_*i*_), where *i* represents the generic index of the training MS images, *f*_*i*_ is the colocalization feature vector extracted from *N*_*pix*_ ROI pixels of the i-th MS image (Supplementary Equation 3) and *Y*_*i*_ represents the tissue section label, with *Y*_*i*_ ∈ {breast, colorectal, ovarian}.

The performance of the models was evaluated by comparing the predicted labels of the validation MS images with their true values *Y*_*−i*_, using the colocalization features *f*_*−i*_, where *‘−i’* denotes the MS image indices that did not belong to the *training set* (Figure 1B). In each round of the cross-validation, prediction accuracy, single class sensitivity and specificity were calculated as prediction performance metrics. The average of the performance metrics was calculated at end of the 10-fold cross validation as a representative metric for the predictive power of the model. A varying number of PLS-DA components *K* = {2, 3, 4, …, 10} were tested.

Since the split of MS images into *training* and *validation sets* was generated randomly, and this might affect the values of the performance metrics, the 10-fold cross-validation was repeated 500 times. In each repetition, the colocalization features were calculated using a different set of random *N*_*pix*_ tissue-related ROI pixels.

The average metrics calculated over the 500 repetitions were considered as representative of the performance associated with the classification parameters. By this approach, the maximum average accuracy was used to determine the optimal number of PLS-DA components *K*^*^ and the optimal number of the randomly selected pixels 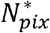.

In order to test whether the prediction performance observed with the *cross-validation set* was due to overfitting, a PLS-DA with the optimal number of components *K*^*^ model was fitted to the colocalization features of the entire *cross-validation set* MS images, then extracted using the optimal number 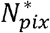 of randomly sampled pixels. After fitting the model, the prediction performance was measured on the external *test set* MS images. Similarly to cross-validation, the entire procedure was repeated 500 times, and the final performance metrics were calculated as the average of the metric values.

### Mean peak intensities as representative features of MS images

A comparison with a spectral mean-based procedure for whole MS images classification was performed. Each MS image was represented by a vector consisting of the mean intensities of the matched peaks, calculated using the tissue-related ROI pixels only.

These vectors were used as features for a PLS-DA supervised model. An identical scheme to that employed with the colocalization features was used to evaluate the best combination of a number of pixels and PLS-DA components. The best mean accuracy obtained with this method was finally compared with that obtained from the colocalization feature-based classifiers.

All the scripts were developed in the R language^45^ and are available at https://github.com/paoloinglese/ion_colocalization.

## RESULTS

The pre-processing workflow applied to the *cross-validation set* resulted in 64 common peaks. When assigned to the 21 external *test set* MS images, it appeared that one sample had one unmatched peak, two samples had two unmatched peaks, and one sample had three unmatched peaks. The small number of unmatched peaks suggested that the common peaks determined by our procedure were generally present in the three classes of tissues. The resulting 64 common peaks were annotated by generating the sum formula from the exact mass measurements and by searching the *m/z* values at the online Metlin database, HMDB and SciFinder^46–48^ (Supplementary Table S4). An *m/*z error threshold 5 ppm was used to accept the putative annotation.

Further structural elucidation was performed using MS/MS experiments via collision-induced dissociation on the LTQ-Discovery MS instrument (Thermo Scientific). For the annotation of metabolites, the MS/MS spectra were matched against spectral libraries from HMDB, NIST and Metlin that were compiled with either authentic standards or theoretical assignment. Identification of metabolites whose MS/MS spectra are not present in the spectral libraries remained challenging, and ions were annotated based on known chemical rule-based fragmentation pattern. For the structural assignment of glycerophospholipids, fragments of the polar head group or the fatty acyl chains were investigated to confirm the annotation proposed by the databases and discriminate isomers.

### Colocalization feature variance converges to zero with a larger sets of pixels

A first analysis aimed to determine the variability of the 2,016 colocalization values due to the randomness of the ROI pixels used to extract the features. The variance of the entire colocalization vector values across the 500 repetitions was calculated for each MS image (Supplementary Methods 1.5). As expected, with an increasing number of pixels, the variances converged to 0. In particular, with a sample size larger than 200 pixels, the colocalization features variance was smaller than 0.01 (Figure 2).

**Figure 2.**
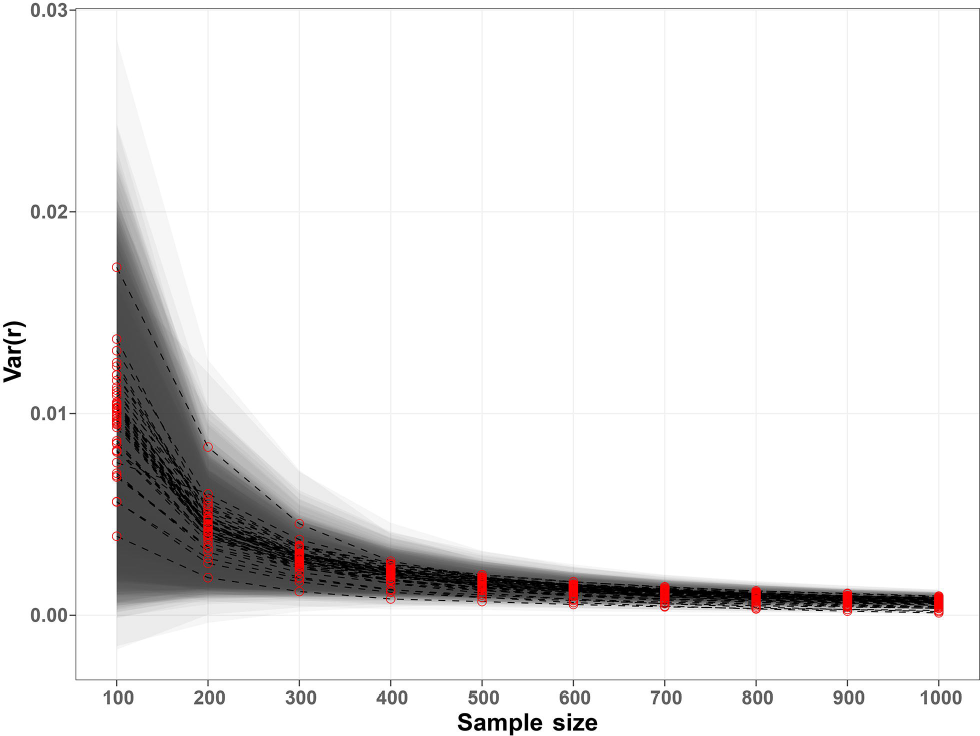
Variance of the colocalization vectors values. The red dots represent the median of each MS image colocalization features calculated over the 500 repetitions using *N*_*p*_ randomly sampled pixels. The bands represent the 1.4826 * MAD distance from the median. The plot shows that the variance of the colocalization features converged to zero with an increasing number of pixels, suggesting that these features were almost independent from the used pixels with samples larger than 300 pixels.

### Performance of colocalization-based models

The 2,016 Spearman’s correlations were used as colocalization features for the PLS-DA models. A number 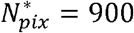 of randomly sampled pixels and K^*^ = 7 PLS-DA components corresponded to the maximum average accuracy across the 500 repeated 10-fold cross-validations (Supplementary Equation 6) (Figure 3), equal to 0.9158±0.0262 (Table 1). A comparison with the accuracies obtained after permuting the MS image labels confirmed that the observed performances were not due to random chance (Supplementary Figure S2). The optimal number of sampled pixels 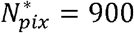 represented a percentage of the tissue-related ROI size varying from 6% to 75% with a median of 21%.

**Table 1.** Tissue classification performance (average ± standard deviation) of the PLS-DA models, using the optimal number of pixels equal to 900, and the optimal number of PLS-DA components equal to 7. The cross-validation results represent the performances of the models on the 10-fold cross-validation, the external test results represent the performances of the prediction of the external test MS images using the optimal parameters identified by the cross-validation.

| Cross-validation |  |  |  |
| --- | --- | --- | --- |
| Accuracy | 0.9158±0.0262 | | |
|  | Breast | Colorectal | Ovarian |
| Sensitivity | 0.9030±0.0385 | 0.9471±0.0195 | 0.9840±0.0308 |
| Specificity | 0.9731±0.0255 | 0.8605±0.0589 | 0.9536±0.0254 |
| External test |  |  |  |
| Accuracy | 0.8059±0.0346 | | |
|  | Breast | Colorectal | Ovarian |
| Sensitivity | 0.6775±0.0481 | - | 0.9771±0.0482 |
| Specificity | 0.9911±0.0310 | - | 0.6775±0.0481 |

**Figure 3.**
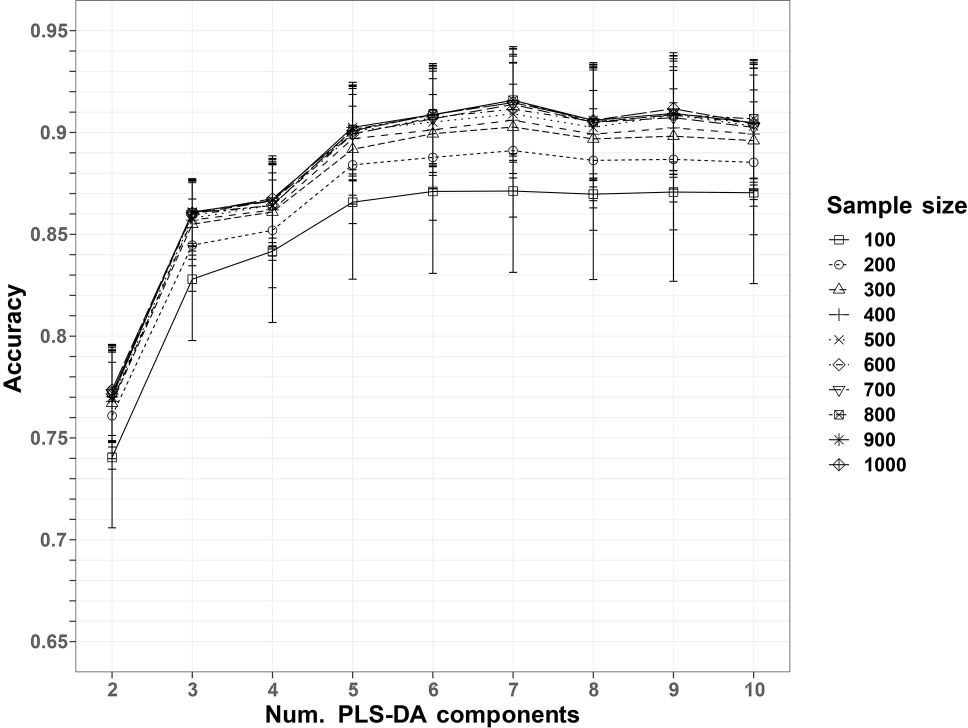
Accuracies of the repeated PLS-DA classifications. The dots represent the average accuracies of the 10-fold cross-validation, repeated 500 times. The error bars represent the standard deviations. The maximum average accuracy corresponds to a sample size of 900 pixels and a number of PLS-DA components equal to 7.

The PLS-DA model trained on the whole *cross-validation set* was capable of predicting the external *test set* labels with an accuracy of 0.8059±0.0346 (Table 1). This result confirmed that the model was adequately capable of distinguishing between tumor types.

### Performance of mean peak intensity-based models

The classification models using the mass spectral intensities as features resulted in lower cross-validation performances compared to the co-localization-based models, with an average and standard deviation accuracy of 0.8138±0.0324 (Supplementary Figure S3).

### Bias analysis of the three datasets

Since the datasets used in this study were obtained from three different studies, we tested for the presence of batch effect that could cause bias in the fitted models.

Two types of batch effects were considered as possibly related with the ion colocalization. Firstly, a systematic variation of the *scatteredness* of the ion images among the three datasets. Secondly, a batch effect represented by a systematic variation in the peak intensities associated with the three datasets.

Given an ion image, its scatteredness was defined as the ratio between the number of pixels in non-zero-intensity connected regions (two pixels are connected if they are 1-nearest-neighbours) and the total number of non-zero-intensity pixels in the tissue-related ROI. This measure was intended to capture the degree of sparsity of the image since more sparse images are characterized by larger fractions of disconnected signal pixels. Being based on ion image correlations, the colocalization features could be therefore affected by a systematic difference of the scatteredness in the three datasets.

In order to determine the presence of a scatteredness batch effect, a univariate Kruskal-Wallis^49^ test was performed on the scatteredness of each ion images of the three datasets MS images. Specifically, for each ion, the value of scatteredness was calculated for all of the 48 MS images, and the test was used to determine whether a significant difference was present between the three datasets. After a Benjamini-Hochberg correction, none of the 64 ion scatterednesses was found different, at a level of significance of 95% (Figure 4).

**Figure 4.**
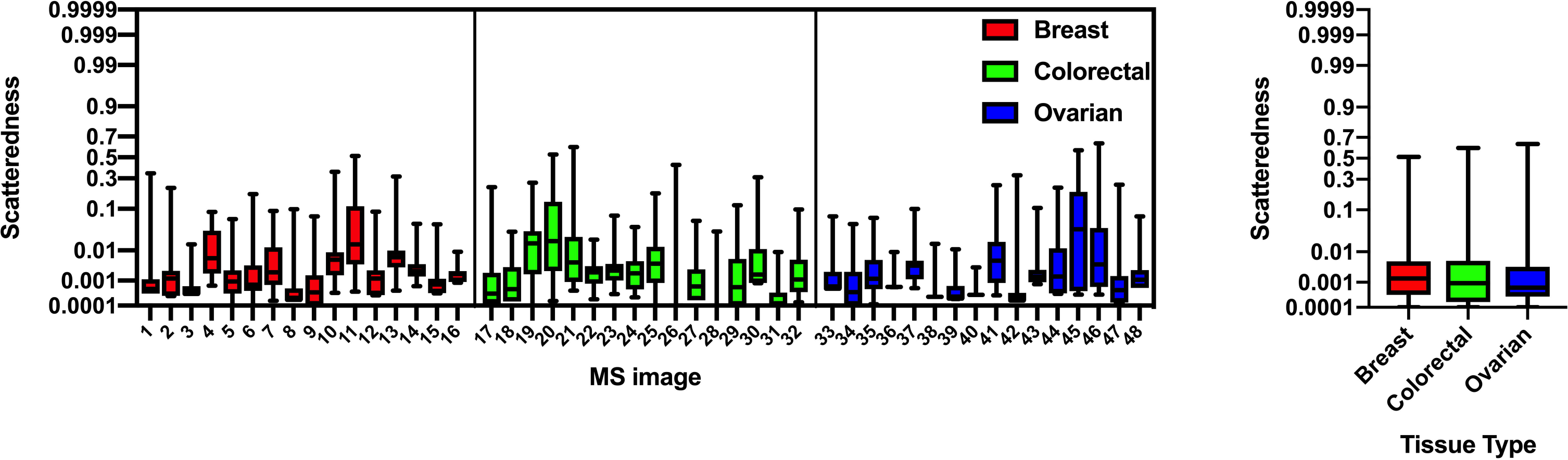
Left: Box plot showing the scatteredness of the ion images of the three sets of MS images. Right: Box plot of the combined scatteredness of the three main tissue types. A Kruskal-Wallis test resulted in no difference between the tissue types, at a level of significance of 95%.

The peak intensity batch effect was tested similarly. All the peak intensities of all the MS images were combined, and a Kruskal-Wallis test was performed to determine if a significant difference was observed between the three datasets. In this case, a significant difference (p < 1e-5) between the three datasets was observed (Figure 5A-B).

**Figure 5.**
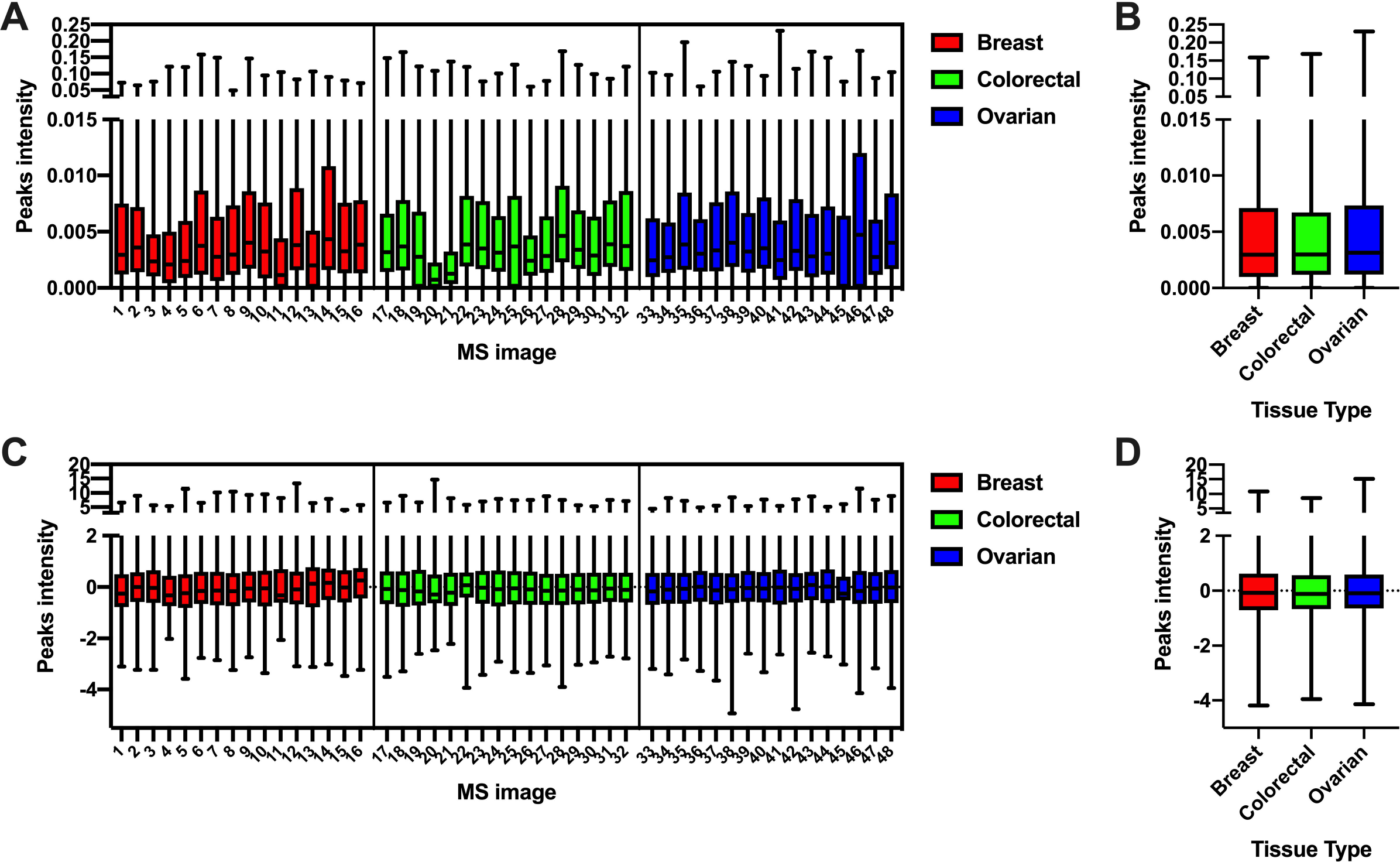
Box plot of the preprocessed peak intensities of the 48 MS images belonging to the cross-validation set (A) and combined in the three main tissue types (B). A Kruskal-Wallis test resulted in a difference in the intensity distribution between the tissue types, at a significance level of 95%. After standardization (C-D), no significant difference was found between the peak intensities of the three tissue types.

In order to test whether this systematic variation was responsible for the performance of the colocalization-based PLS-DA models, we repeated the classification using batch-effect corrected peak intensities. Peak intensities were standardized within each MS image. Specifically, for each MS image, intensities of individual peaks were centered using their mean values and scaled dividing by their standard deviation. After applying the scaling, no significant difference was observed between the three datasets (Figure 5C-D).

Using the optimal parameters, 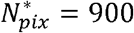 and K^*^ = 7, the cross-validation accuracy with the colocalization features of the standardized intensities was almost identical to that observed using the unscaled intensities, resulting equal to 0.9157±0.0269. This result confirmed that the colocalization features were insensitive to the relative differences of peak intensities between the three datasets.

### Colocalization features are less sensitive to batch effect

In order to test the robustness of the colocalization features to peaks intensity batch effects, we performed a series of tests with the application of simulated intensity offsets. In each set of experiments, the MS images from only one tissue type were considered. Firstly, all the peak intensities were standardized (as described in the previous section). Afterwards, a numeric offset was applied to the peak intensities in half of the 16 MS images. This offset was different for each pixel and peak (to simulate also small variations within the MS images) and randomly sampled from a Normal distribution with a varying mean in 0.1, 0.5, 1, 2 and standard deviation equal to 0.1.

The aim of this approach is to determine whether a PLS-DA model was capable of distinguishing MS images with or without the applied offset. A good accuracy would then imply that the features are sensitive to the presence of systematic variations.

A classification procedure similar to that described in the previous sections was applied, with the only difference of using a leave-one-out cross-validation, due to the smaller number of tested samples (16 MS images per tissue type).

Performance was calculated using the colocalization features and the mean peak intensity features for comparison purposes.

As expected, the results showed that colocalization features are much less sensitive to systematic variations than the mean peaks intensity features, with the latter giving an accuracy of 1.0 with offsets sampled from a normal distribution with mean equal to 0.1. In contrast, the accuracy of the colocalization features remained almost constant (Figure 6).

**Figure 6.**
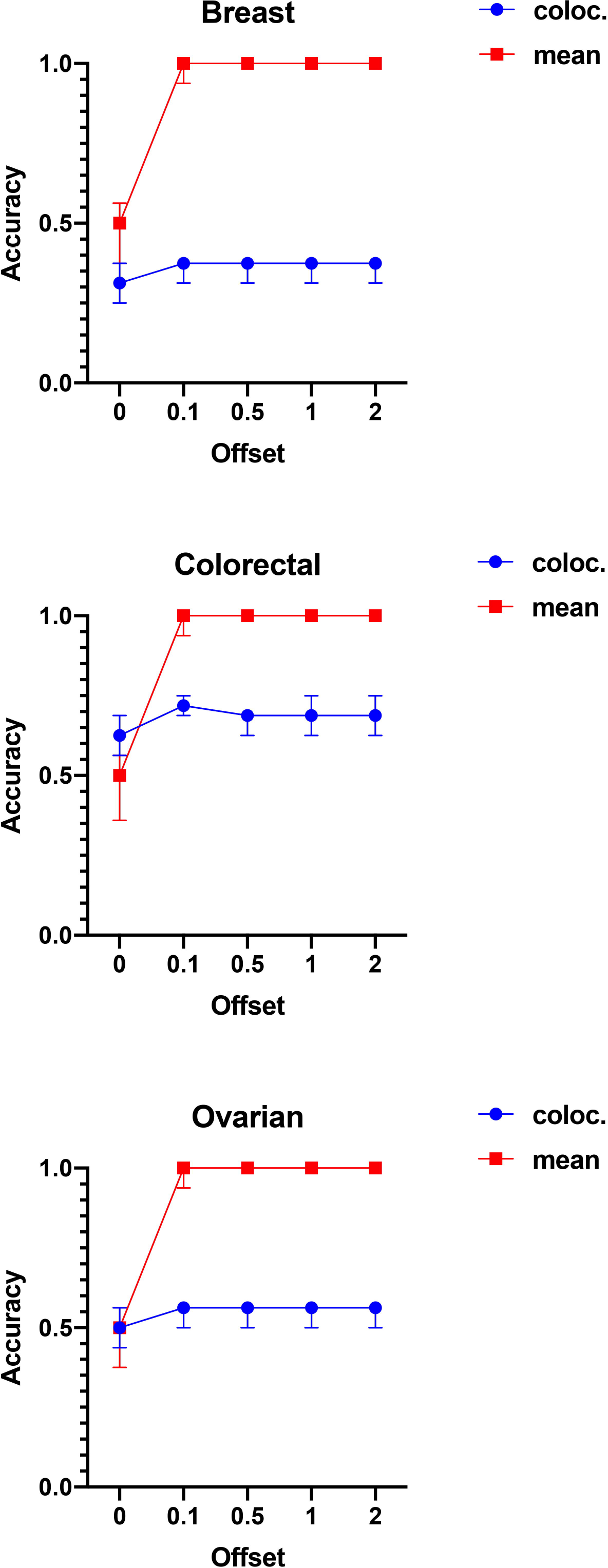
Best accuracies for the discrimination between MS images with or without applied offset. The results show that the mean peaks-intensities were significantly more sensitive to the applied offsets, allowing the PLS-DA models to predict them correctly. On the contrary, the PLS-DA models trained on the colocalization features performed almost identically irrespective of the offset distribution, confirming that these features are more robust to systematic variations of the peak intensities.

### Univariate test of colocalization features and visualization

A multiple pairwise univariate Kruskal-Wallis test was employed to determine the colocalization features that were significantly different between the three tumor types. The features selected with more than a 95% significance level (after Benjamini-Hochberg correction, with a number of tests equal to 3 × 500) were further investigated to identify the most informative colocalization patterns for the discrimination of the three tumor classes. It was observed that, across the 500 repetitions, the number of significantly different correlations were stable (Supplementary Figure S4). The features that were significant in the 95% of the 500 repetitions were used for the following analysis. Their overall significance was calculated as the average adjusted p-values across the 500 repetitions, and the features were sorted by their overall significance (increasing order).

The ten most significant correlations are reported in Supplementary Table S6 together with the elemental composition corresponding to the absolute *m/z* differences between the ion pairs associated with the selected correlations that can provide information about the possible chemical interactions.

The spatial localization of these ions could be investigated by plotting the relative abundance images of the spectral peaks involved in the selected correlations (the spatial distribution of the most significant correlation is reported in Supplementary Figure S5). These images confirmed that these ions were localized in tissue-related areas and did not involve off-tissue regions of the images. In order to compare the spatial distributions of the ion pairs involved in the selected correlations with the corresponding optical images of the H&E stained tissue, we segmented the MS image into two clusters. The main cluster (represented by the yellow pixels in the Supplementary Figure S6) represented the pixels where both correlated ions had a higher relative abundance than their average value. In this way, it resulted that the ions associated with the most significantly different correlations in the breast/colorectal and colorectal/ovarian contrasts were mainly localized in the tumor region, whereas in the breast/ovarian contrast they were mainly localized in the connective tissue (Figure 7).

**Figure 7.**
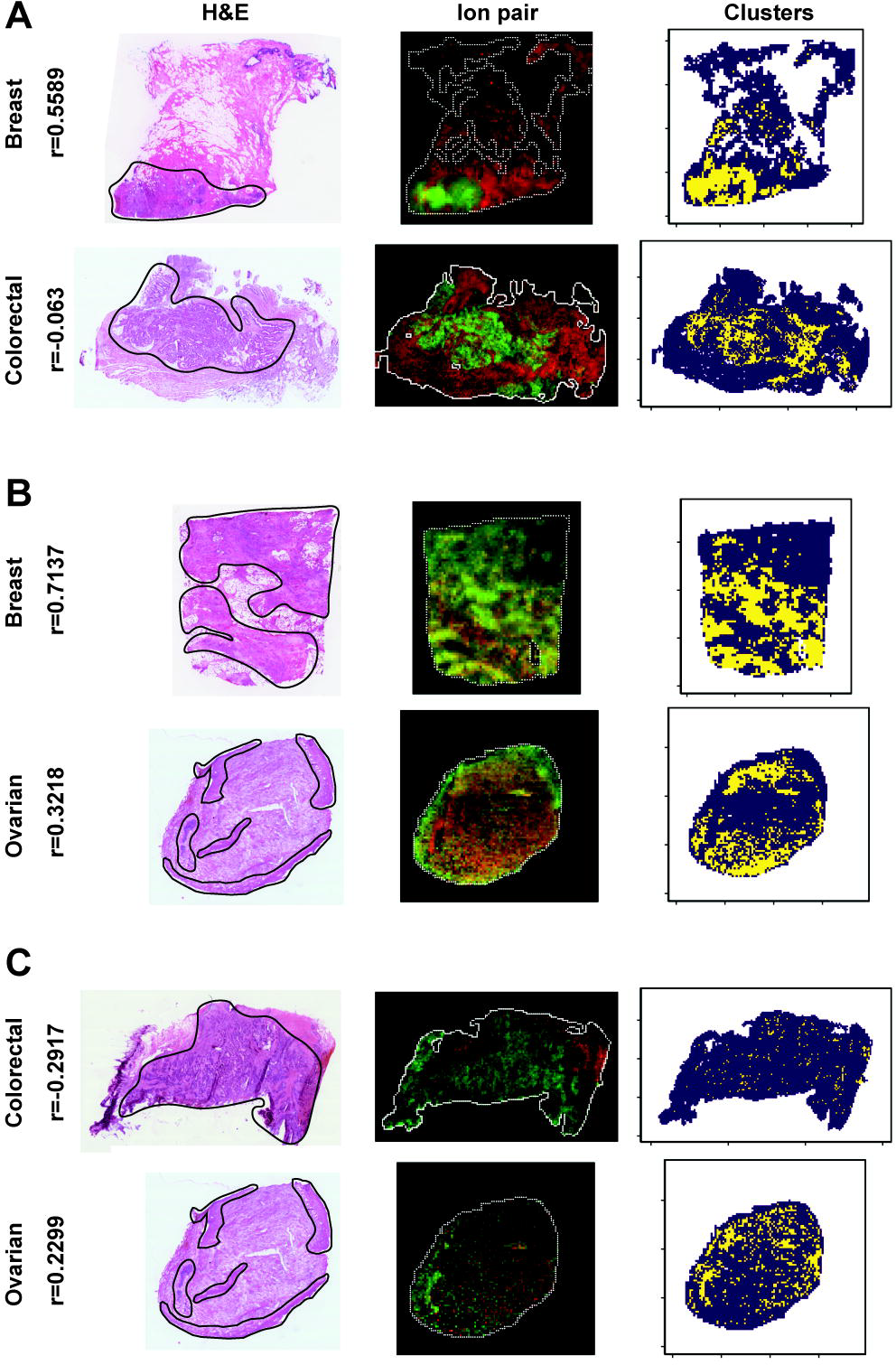
Example of the spatial patterns associated with the most significant correlation in the three contrasts. The shown DESI-MS images had the closest selected correlation values to the median within the tissue group. The selected correlations ions are mainly localized in the tumor tissue (delineated by a black line in the H&E optical images) for the breast/colorectal (A) and colorectal/ovarian (C) contrasts, whereas it is mainly localized in the connective tissue in the breast/ovarian (B) contrast, as revealed by the clusters (third column). The yellow pixels represent the regions of the tissue where the two ions have a relative abundance higher than the average value within the MS image. In the second column, the ion pairs abundances (scaled in [0, 1]) are represented as red and green channels of an RGB image. The Spearman’s correlation values (r) are reported on the left side.

For each repetition, the selected correlation features associated with the three pair-wise comparisons were averaged across the MS images of each tissue class as representative values of their group. Afterwards, a circle plot of the average of the repeated values revealed the different colocalization patterns between each pair of tissue class (Figure 8).

**Figure 8.**
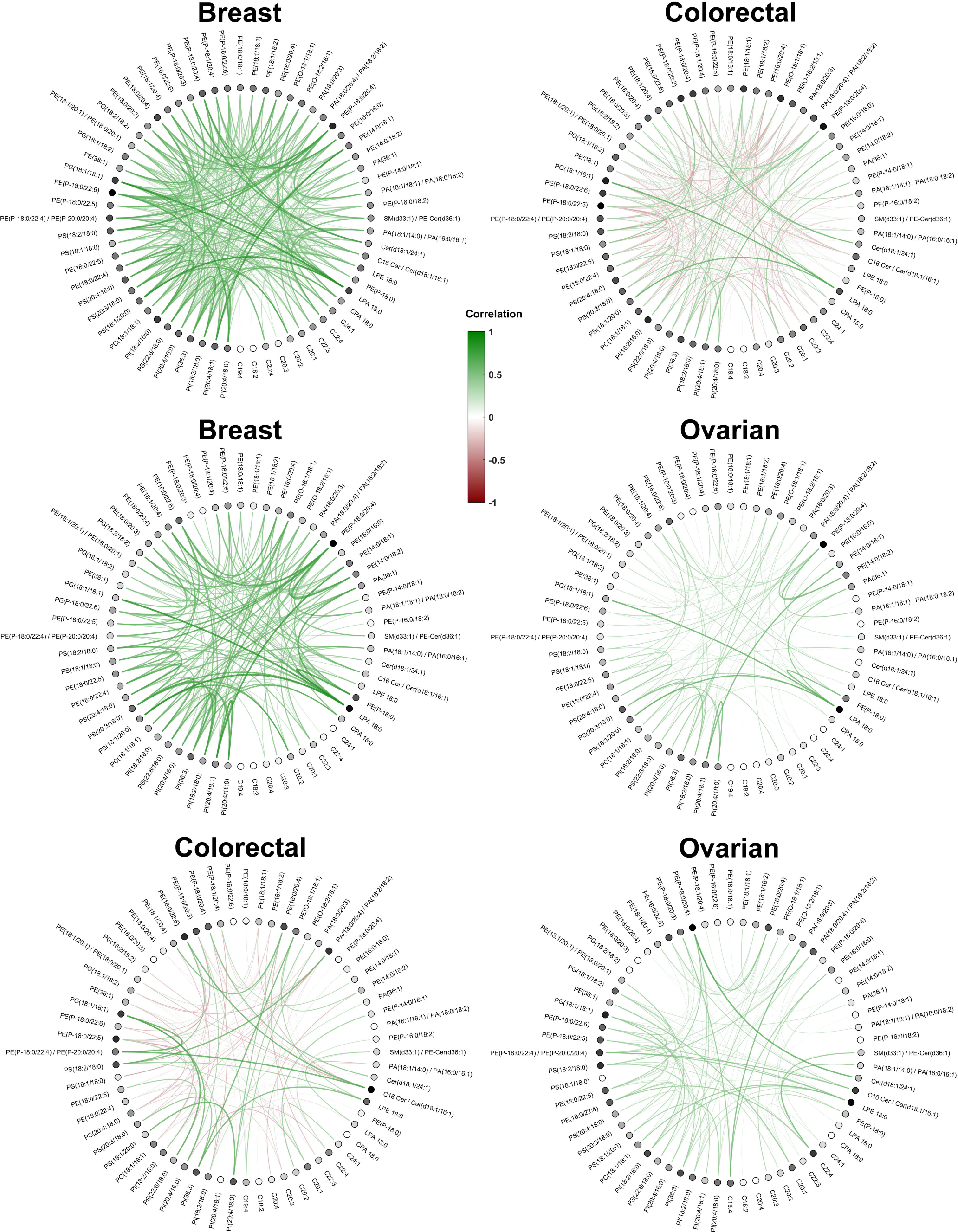
Circle plots representing the correlation features that were significantly different in the three pairwise contrasts. Each row represents the significant correlations averaged across the MS images of the tissue class and the 500 repetitions. Positive correlations are plotted in green, whereas negative correlations are colored in red. Each dot represents the molecule ion involved in the colocalization feature, with a darker color for nodes that were involved in higher absolute average values of correlations. The graphs illustrate clear differences in colocalization patterns between the three tumor types.

A graph representing the significant correlations was determined from each pairwise tumor comparison. The average correlations calculated across the 500 repetitions within the same tumor class were used as the edges of a circle graph, whereas the peak *m/z* corresponding to the ion pairs were assigned to the nodes (Figure). This graph revealed the different colocalization patterns associated with the three types of tumors. Following the assumption behind the workflow presented, these patterns can be used to generate hypotheses based on the interactions that distinguish the biochemical characteristics of the different cancer types.

## DISCUSSION

Mass spectrometry imaging is gaining increased interest as a technology capable of mapping the spatial distribution of the molecules of interest in two or three-dimensional clinical samples. However, the quantitative analysis of imaging data is still limited due to difficulties controlling factors such as matrix effects. Ion colocalization analysis, however, can represent a more robust alternative towards more quantitative analysis of this kind of data. Colocalization features were insensitive to peaks intensity batch effects, as seen from experiments on the simulated intensity offsets. This result highlights one of the main advantages of the colocalization feature representation, allowing the comparison between MS images obtained from different studies.

Biologically, the workflow exploits the assumption that local metabolic pathways are expressed through colocalization patterns of the molecules involved in the mechanisms. The two strongest types of biological correlations are (i) when the two correlated species are the substrate and the product of the same enzyme, or (ii) when both are products or both are substrates of the same enzyme/transporter. Typical examples for both cases are shown in Supplementary Table S5. which summarizes the strongest correlations for the individual datasets. The strongest correlation (0.92) in the entire dataset was found to be between PA(36:1) and PS(18:0/18:1), one is a potential precursor assuming formation via phospholipase D activity. Overexpression of phospholipase D is one of the common hallmarks of epithelial cancers, activating anti-apoptotic pathways^50^. Examples for parallel biosynthesis of species showing strong correlations include PG(18:1/18:1) and PG(18:1/18:2) which are the products of the same phosphoglycolate phosphatase.

Specifically, we hypothesize that different classes of samples (different tumor types in this case) are characterized by diverse metabolic pathways. This hypothesis was corroborated by the high predictive power of the supervised models. Similar performance was observed in an external *test set*, confirming the robustness of the models. The method outperformed the classification performance where the mean spectral intensities were used as features.

Interestingly, we observed a small variance for the colocalization features, with a value of about 0.01 for samples of 300 pixels. Therefore, most of the variation observed in the classification performance metrics was due to the different split into *training/validation sets* in each repetition. In other words, we did not observe sub-regions characterized by different colocalization patterns (e.g. two ions were positively correlated in a portion of the tissue and anti-correlated in another sub-region) inside the same tissue section. This can be a result due either to underlying stability of the detected biochemical mechanisms or to the technical limitations of the analytical technique employed to generate the MS image.

Additionally, the colocalization patterns can be used for data-driven hypothesis generation, suggesting possible local molecular mechanisms characterizing the samples of interest. The visualization of the most significantly correlated ions can help to determine regions of interest for further experiments, through data integration with transcriptomics or simply by inspection of the stained samples for histopathological validation, in the case of tissue sections. When represented as a graph, the significant correlations reveal patterns that could be used as a part of the hypothesis generation process.

Both positive and negative correlations were considered as feature values for representing the MS images. Positive correlations may indicate the presence of one or multiple mechanisms locally involving the detected molecules, whereas negative correlations may be seen as reflecting competitive processes like COX1/2, in which arachidonic acid is used as a common precursor whereas prostaglandins PG H2 and PG G2 are produced depending on the COX expression^51^. However, MSI data alone cannot provide conclusive evidence of such mechanisms.

Another limitation of the method is characterized by the representativeness of the specimens studied. A single tissue section may be not representative of the complex heterogeneity of the tumor. For this reason, multiple sections or three-dimensional specimens may give a more robust representation of the molecular composition of the analyzed samples^15^.

The dimensionality of the colocalization features defined in the presented work is *O*(|*v*|^2^) where |*v*| represents the number of spectral peaks. For this reason, when a large number of spectral peaks are available, then the colocalization feature vectors dimensionality can become computationally intractable. In such situations, feature selection methods (e.g. Random Forest variable importance) may help to reduce the number of features used to train the classification models.

In the future, various similarity measures will be tested for the quantification of the degree of ion colocalization in cancer tissue and MSI data from different sources. Additionally, biologically-related hypotheses generated by the ion colocalization patterns will be thoroughly investigated through extensive experimental validation, involving transcriptomics and immunohistochemistry measurements.

## ACKNOWLEDGEMENTS

We would like to thank Dr Luisa Doria, Dr Sabine Guenther, Ms Anna Mroz, Dr Nicole Strittmatter for sharing the datasets used in their publications.

This work was supported by Cancer Research UK Grand Challenge Program (Project title: ‘A Complete Cartography Through Multiscale Molecular Imaging’). The Department of Surgery and Cancer is funded by grants and infrastructure support by the National Institute for Health Research (NIHR) Imperial Biomedical Research Centre (BRC). GC is supported by the National Institute for Health Research (NIHR) Imperial Biomedical Research Centre (BRC). Funding for PP was provided by a European Research Council Consolidator Grant (617896). The views expressed in this publication are those of the author(s) and not necessarily those of the NHS, the National Institute for Health Research or the Department of Health.

