## Supplementary for "Colocalization features for classification of tumors using desorption electrospray ionization mass spectrometry imaging"

2 Table of Contents

### 1 SUPPLEMENTARY METHODS

#### 1.1 Co-localization features extraction

Let $\boldsymbol{I}_{t}$ be a set of mass spectral intensities associated with a tissue section denoted by the subscript $t$. Given the spatial dimensions $n_{x}$ and $n_{y}$of $t$ in number of pixels, and the *m/z* space dimension $n_{mz}$, then $\boldsymbol{I}_{t}$ can be defined as a 3-dimensional matrix

$$\boldsymbol{I}_{t}=\left( y_{ijk} \right), i\in\left\{ 1,\ldots,n_{x} \right\}, j\in\left\{ 1,\ldots, n_{y} \right\},k \in\left\{ 1,\ldots, n_{mz} \right\}$$

(1)

where $y_{ijk}\in\mathbb{R}_{+}$ represents the $k$-th spectral peak intensity in the spatial location $\left( i,j \right)$. Using this definition, the 2-dimensional sections of $\boldsymbol{I}_{t}$ corresponding to a fixed value of $k$, denoted $y_{..k}$, can be interpreted as images representing of the spatial distribution of the $k$-th spectral peak.

Since $\boldsymbol{I}_{t}$ may contain spatial regions unrelated with the sample-of-interest, the subset of pixels ${ROI}_{t}=\left( i,j \right)\subseteq\left\{ 1,\ldots,n_{x} \right\}\times\left\{ 1,\ldots,n_{y} \right\}$ associated with the tissue section are extracted by applying the ‘kmeans2’ method from the SPUTNIK package for R (Inglese, Correia, *et al.*, 2018) (Supplementary Table S3). The tissue-related set of spectral intensities are denoted as

$$\boldsymbol{I}_{t}^{ROI}=\left( y_{ijk} \right), \left( i,j \right)\in{ROI}_{t},k \in\left\{ 1,\ldots, n_{mz} \right\}$$

(2)

A co-localization features vector of $\boldsymbol{I}_{t}$ is therefore defined as the $s$-tuple

$$\boldsymbol{f}_{t}=\left( f_{1}^{(t)}, f_{2}^{(t)}, f_{3}^{(t)},\ldots, f_{s}^{(t)} \right)$$

(3)

consisting of the Spearman’s correlations between all the pairs of vectorized image pixels $y_{i^{'}j^{'}a}$, $y_{i^{'}j^{'}b}\subset\boldsymbol{I}_{t}^{ROI}$,

$$\begin{matrix} \begin{matrix} \\ f_{p}^{(t)}=Spearman\left( Vectorize\left( y_{i^{'}j^{'}a} \right), Vectorize\left( y_{i^{'}j^{'}b} \right) \right), \end{matrix} \\ p\in\left\{ 1, \ldots,s \right\}, a, b \in\left\{ 1,\ldots,n_{mz} \right\}, \\ i^{'}j^{'}\mathcal{\in R\subseteq}{ROI}_{t} \end{matrix}$$

(4)

such that $a>b$ (or, equivalently, $a<b$). *Vectorize* is a function that concatenates the 2-dimensional spectral peak intensities column-wise, and $\mathcal{R}$ represents a set of randomly sampled ROI pixels, with $\left| \mathcal{R} \right|=N_{pix}$.

*Spearman* is the Spearman’s rank correlation, defined as the Pearson’s correlation between the ranked variables

$$Spearman\left( \boldsymbol{x},\boldsymbol{y} \right)=\frac{\text{cov}\left( rank\left( \boldsymbol{x} \right), rank(\boldsymbol{y}) \right)}{\text{st.dev}\left( rank\left( \boldsymbol{x} \right) \right)\text{×st.dev}\left( rank\left( \boldsymbol{y} \right) \right)}$$

(5)

The correlations were selected to avoid duplicated values, due to its symmetricity under the exchange of the *m/z* indices.

In this way, the vector $\boldsymbol{f}_{t}$ consists of $s=n_{mz}\times(n_{mz}-1)/2$ elements belonging to the interval $\left[ -1,1 \right]$. The asymptotic *t* approximation (Hollander and Wolfe, 1999) is applied to determine the significance of each Spearman’s correlation $f_{p}^{(t)}$. The image correlations associated with a Benjamini-Hochberg corrected p-value (number of tests equal to $s$) larger than 0.05 are set equal to 0.

#### 1.2 Cross-validation scheme

Before fitting the supervised models, the co-localization features are extracted using a randomly sampled number of pixels $N_{pix}$ belonging to the tissue-related ROI.

The 48 MS images belonging to the *cross-validation set* are split into 10 disjoint subsets of almost equal size (their cardinalities must sum to 48) that will represent the validation sets of the 10 rounds of the cross-validation. The co-localization features and the tissue labels of the MS images that do not belong to the $k$-th validation set represent the training set used to train a PLS-DA model. Therefore, the fitted model is used to predict the tissue class of the MS images belonging to the $k$-th validation set. The performance metrics are calculated comparing the true and predicted tissue labels after each round of the cross-validation. At the end of the procedure, the performances are summarized by the average values of the metrics calculated over the 10 rounds. Before fitting the PLS-DA model, all the constant co-localization features in the training set are removed from both training and validation set.

The cross-validation is repeated with a PLS-DA model fitted using a varying number of components in the range of 2 to 10.

The number of the pixels $N_{pix}$ used to extract the co-localization features vary from 100 to 1000 in steps of 100 pixels.

To estimate the effect of the randomized selection of the tissue-related pixels used for the extraction of the co-localization features and the split of the 48 MS images into training and validation set, the entire procedure is repeated 500 times.

The final performances of the PLS-MODELS are summarized by averaging the metrics values of the repeated cross validations for a fixed number of PLS-DA components and number of pixels $N_{pix}$ used to extract the co-localization features.

The optimal number of PLS-DA components $K^{*}$ and the optimal number of sampled pixels $N_{pix}^{*}$ are determined by

$$\underset{\begin{matrix} K\in\left\{ 2,\ldots,20 \right\} \\ N_{pixels}\in\left\{ 100,200,\ldots,1000 \right\} \end{matrix}}{\text{arg }\max} \left( \frac{1}{500}\sum_{i=1}^{500} \left\{ {accuracy}_{i}^{(\text{10-fold})}\left( N_{pix},K \right) \right\} \right)$$

(6)

where ${accuracy}_{i}^{(\text{10-fold})}\left( N_{pix},K \right)$ represents the summary accuracy value of the $i$-th repetition of the 10-fold cross-validation using $N_{pix}$ pixels to extract the co-localization features, and $K$ PLS-DA components.

#### 1.3 External test performance

A PLS-DA model with $K^{*}$ components, fitted on the co-localization features extracted from $N_{pix}^{*}$ (Equation 5) of all the 48 MS images belonging to the *cross-validation set*, is used to predict the labels of the external *test set* MS images.

The procedure is repeated 500 times, with different randomly sampled $N_{pix}^{*}$ pixels.

The final performance metrics are summarized by averaging their values over 500 repetitions.

#### 1.4 Classification using spectral intensity features

In order to compare the performances of the co-localization features with the standard spectral peak intensities, an analogous classification scheme (Supplementary Methods 1.2) was applied to the pixel mass spectral vectors.

The mass spectral intensities vectors associated with the same randomly sampled pixels used to calculate the co-localization features of the training MS images were used as training features for a PLS-DA model.

The performances of the models were measured comparing the true and predicted tissue labels of the test MS images pixels spectra. After the rounds of the 10-fold cross-validation, the performance metrics were summarized by averaging over the 10 rounds.

The entire procedure followed the identical scheme used with the co-localization features.

#### 1.5 Co-localization features variance estimation

Let $\boldsymbol{I}_{t}$ be the generic MS image belonging to the *cross-validation set*, with $t\in\left\{ 1, 2, \ldots, 48 \right\}$. Let $\boldsymbol{f}_{t}^{(r)}=\left( f_{1}^{(t)(r)}, f_{2}^{(t)(r)},\ldots,f_{s}^{(t)(r)} \right)$ represent the its co-localization feature vector (Equation 3) extracted in the repetition $r\in\left\{ 1, 2, \ldots, 500 \right\}$.

The variance of the co-localization vector estimated over the 500 repetitions is the vector of the variances of the co-localization features

$$Var\left( \boldsymbol{f}_{t} \right)=\left( Var\left( f_{1}^{\left( t \right)(.)} \right),Var\left( f_{2}^{\left( t \right)(.)} \right),\ldots,Var\left( f_{s}^{\left( t \right)(.)} \right) \right)$$

(7)

The median of the components of $Var\left( \boldsymbol{f}_{t} \right)$ and its median absolute deviation (MAD) are reported in Figure 2 of the main text.

### 2 SUPPLEMENTARY DATA

Figure S 1 – Example of tissue-related ROI. K-Means clustering with a number of clusters equal to 4 is applied to the MSI intensity matrix. The clusters that are not localized in the off-tissue area (corners used as reference) are merged into the final tissue-related ROI. A comparison with the first principal component scores (PC1) image and the optical image of the H&E stained tissue shows the close correspondence between the ROI and the area occupied by the tissue.

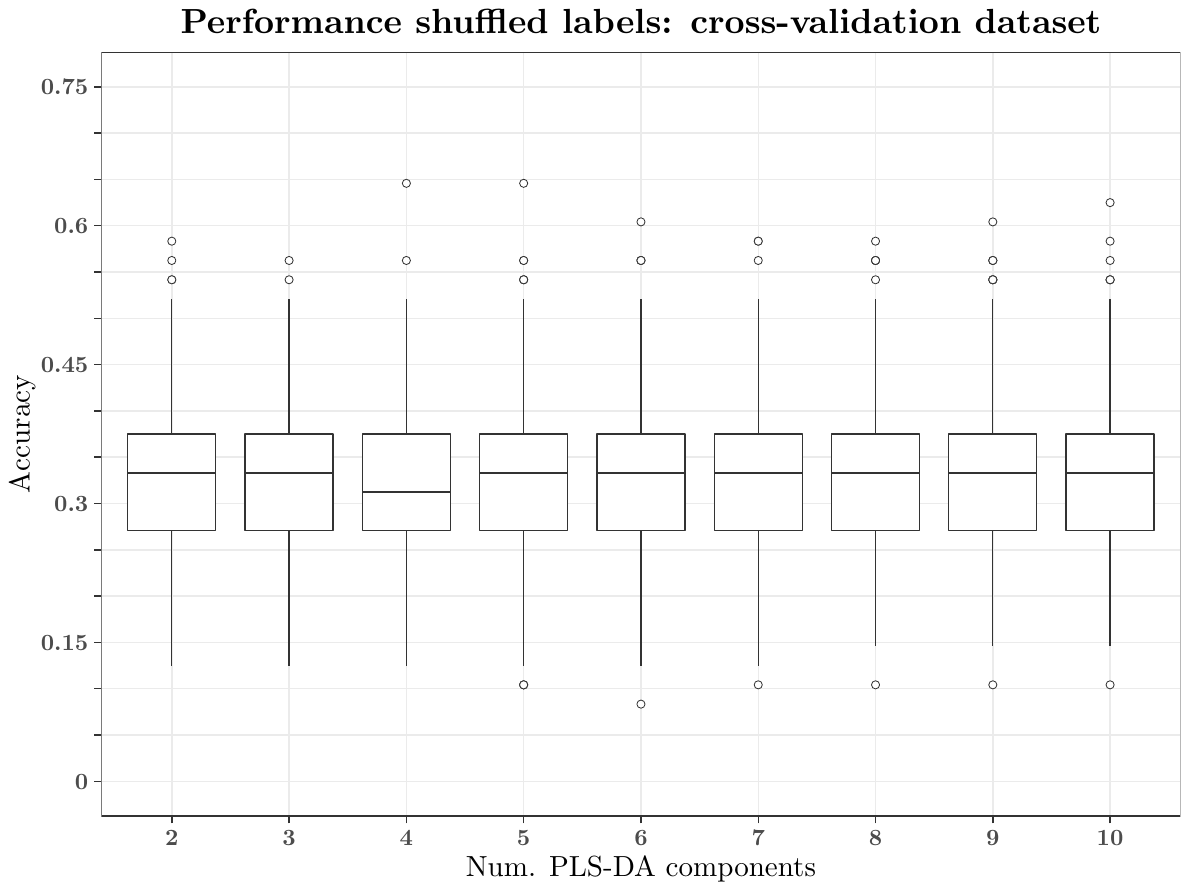

Figure S 2 – Performances of the PLS-DA models with shuffled tissue labels. The bar plots represent the accuracy values of the 500 repetitions using the optimal number of randomly sampled pixels equal to 900, and PLS-DA components varying from 2 to 10. The accuracy values confirm that the observed performances in the original PLS-DA model are not due to random chance.

Figure S 3 - Performances of the PLS-DA models on the cross-validation set, using the mean peak intensities as features.

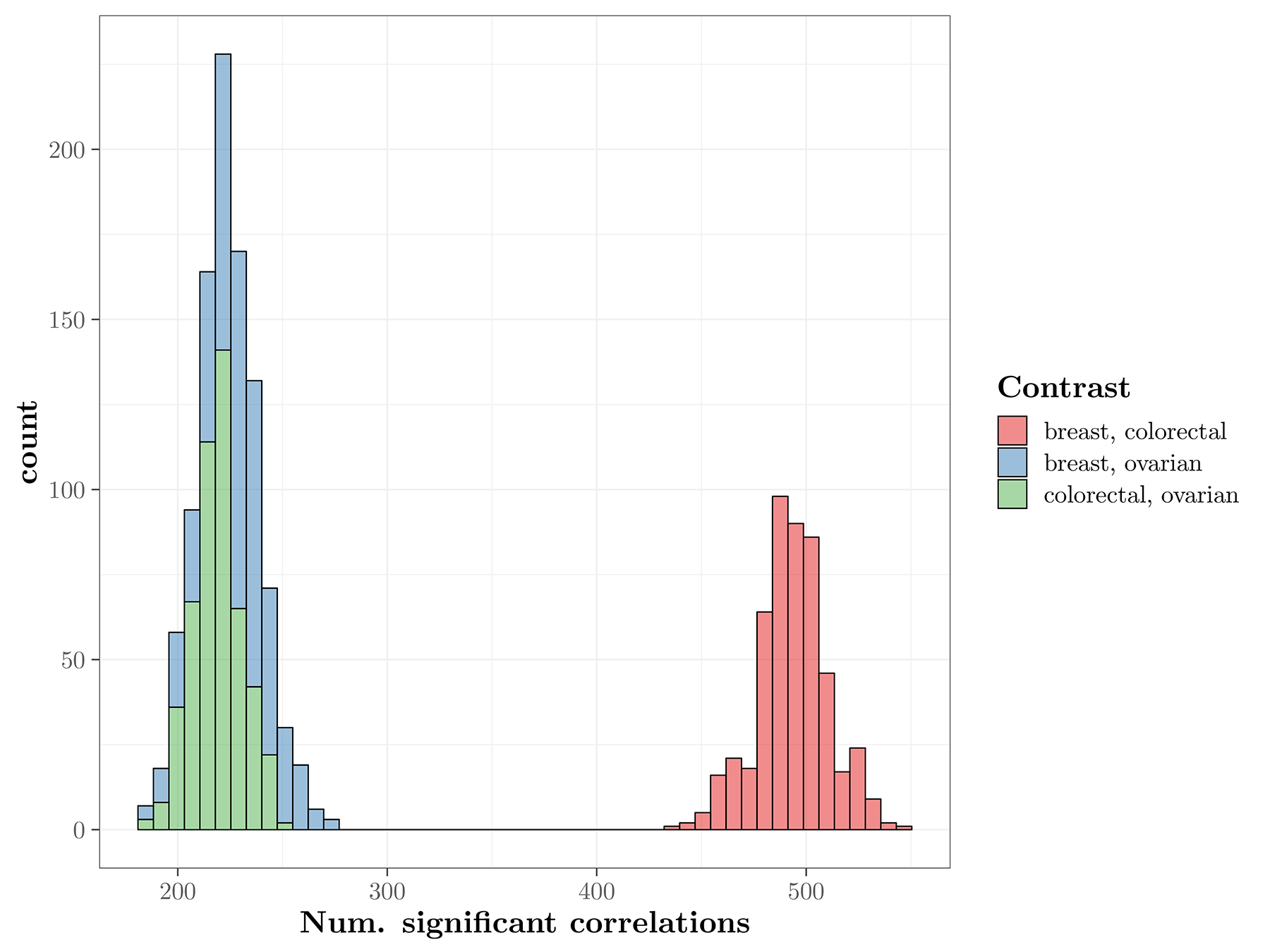

Figure S 4 - Histogram representing the number of significant correlations for the three contrasts, in the 500 repetitions.

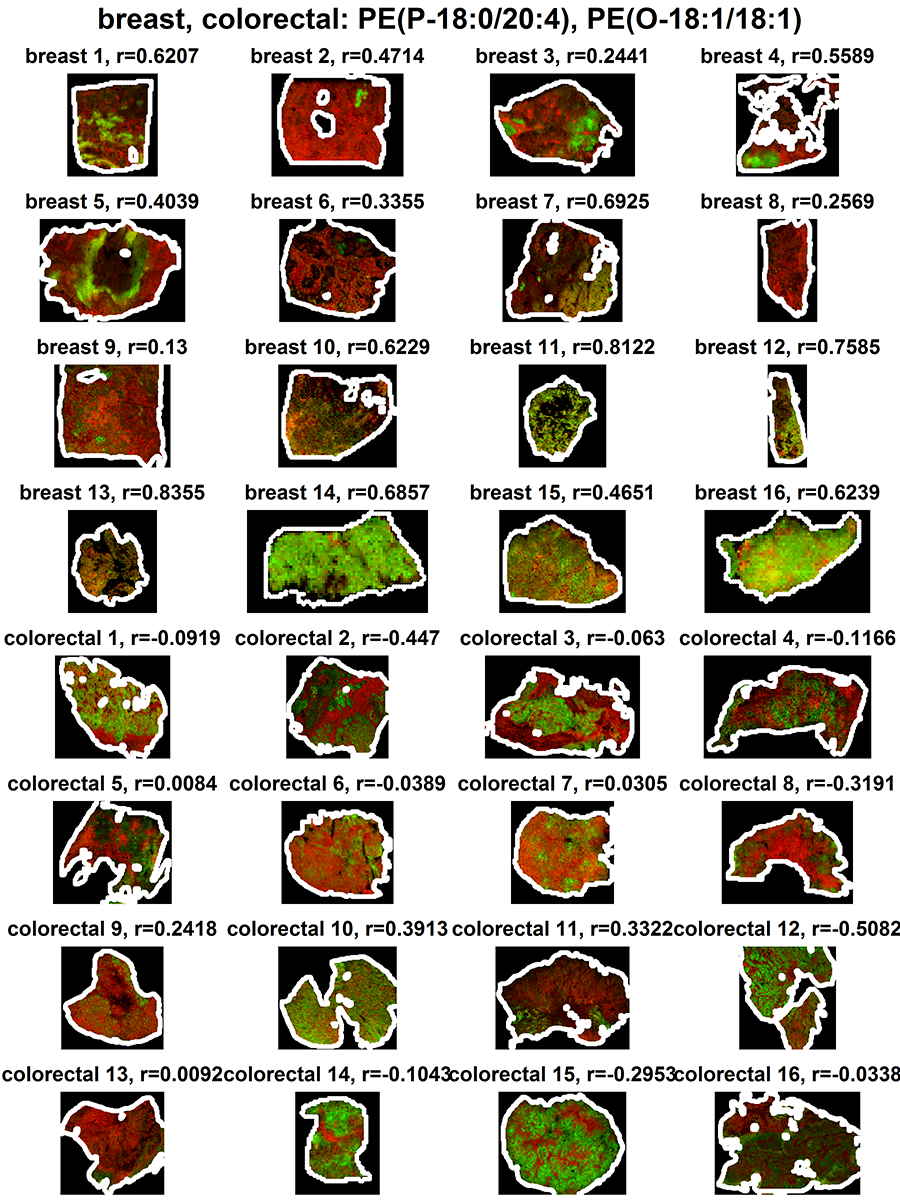

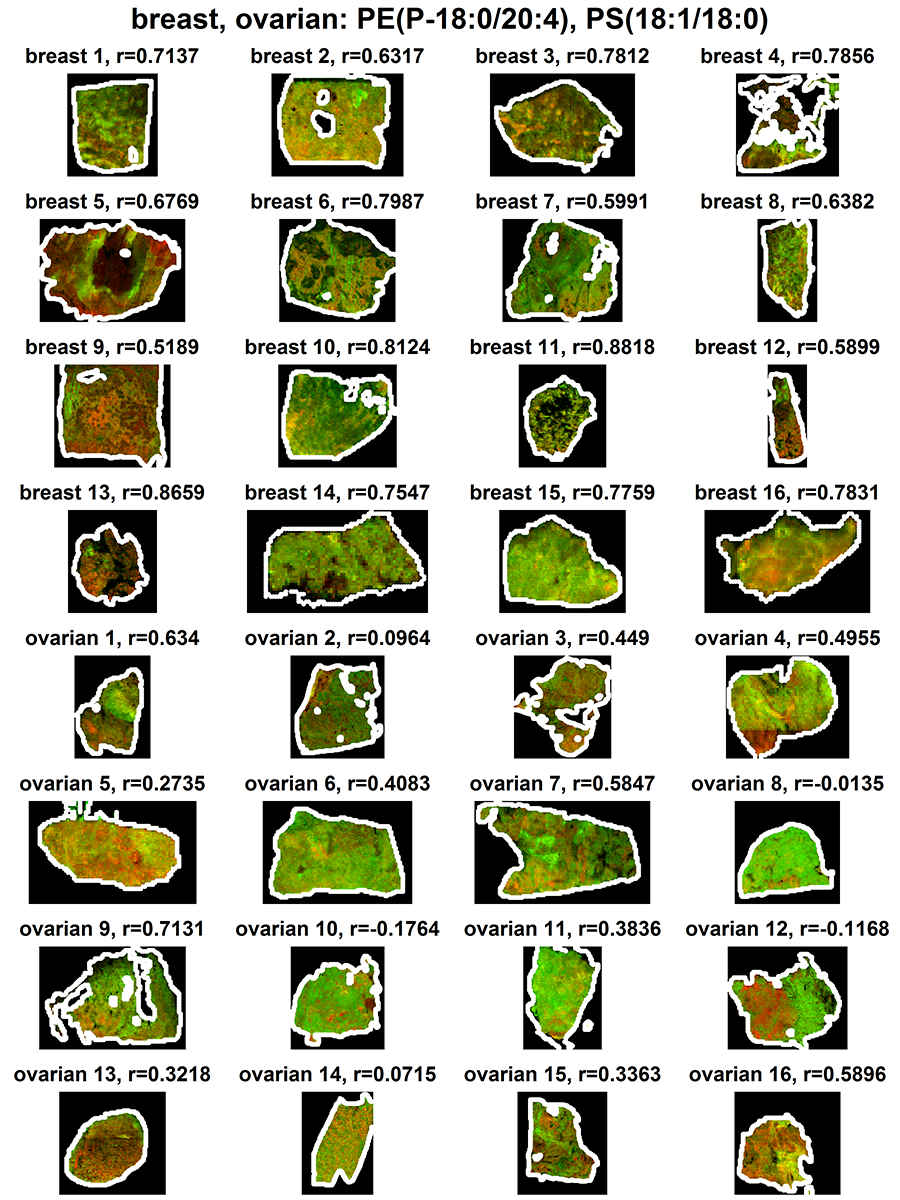

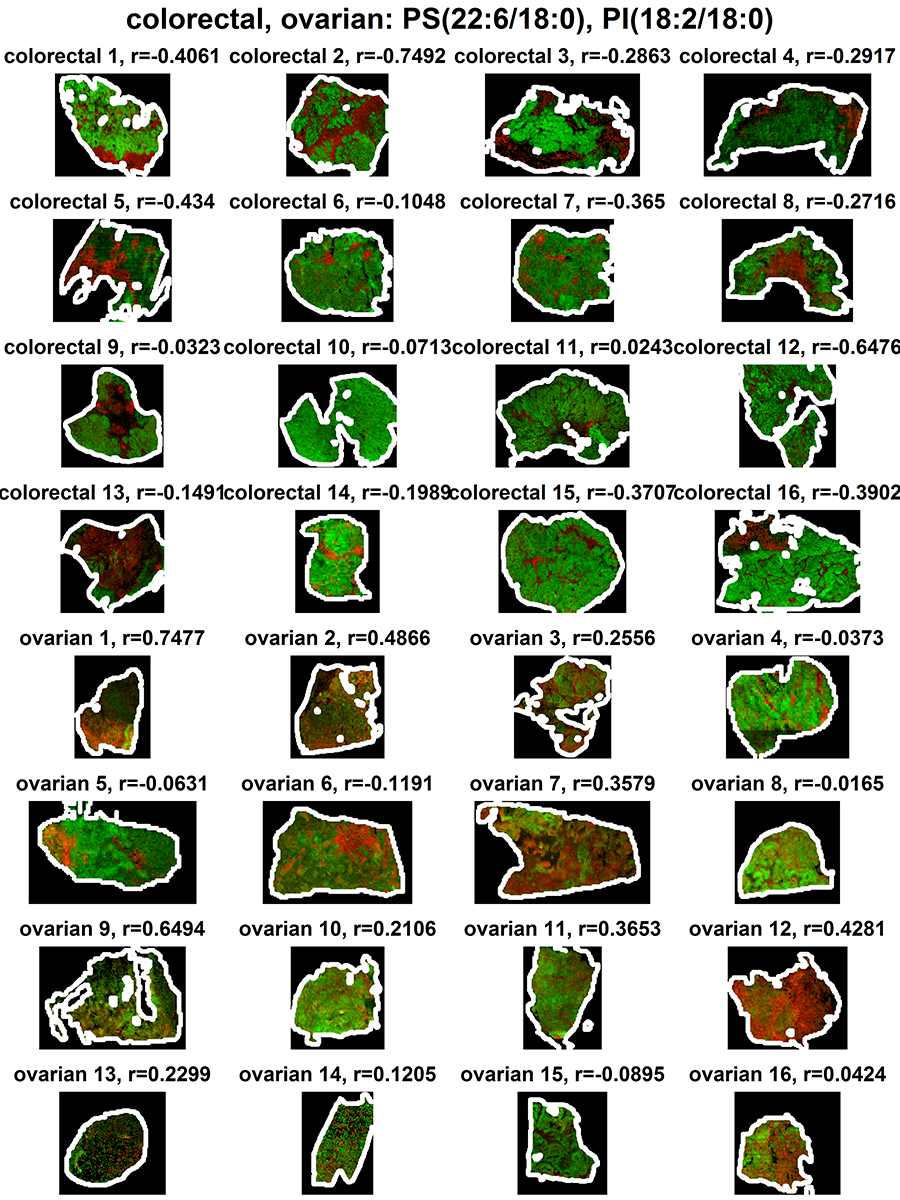

Figure S 5 – Spatial distribution of the spectral peaks associated with the most significantly different co-localization feature in the three contrasts. The relative abundances, scaled in the interval [0, 1], are represented as the red and green channels of an RGB image. The value of the correlation is reported on top of each image.

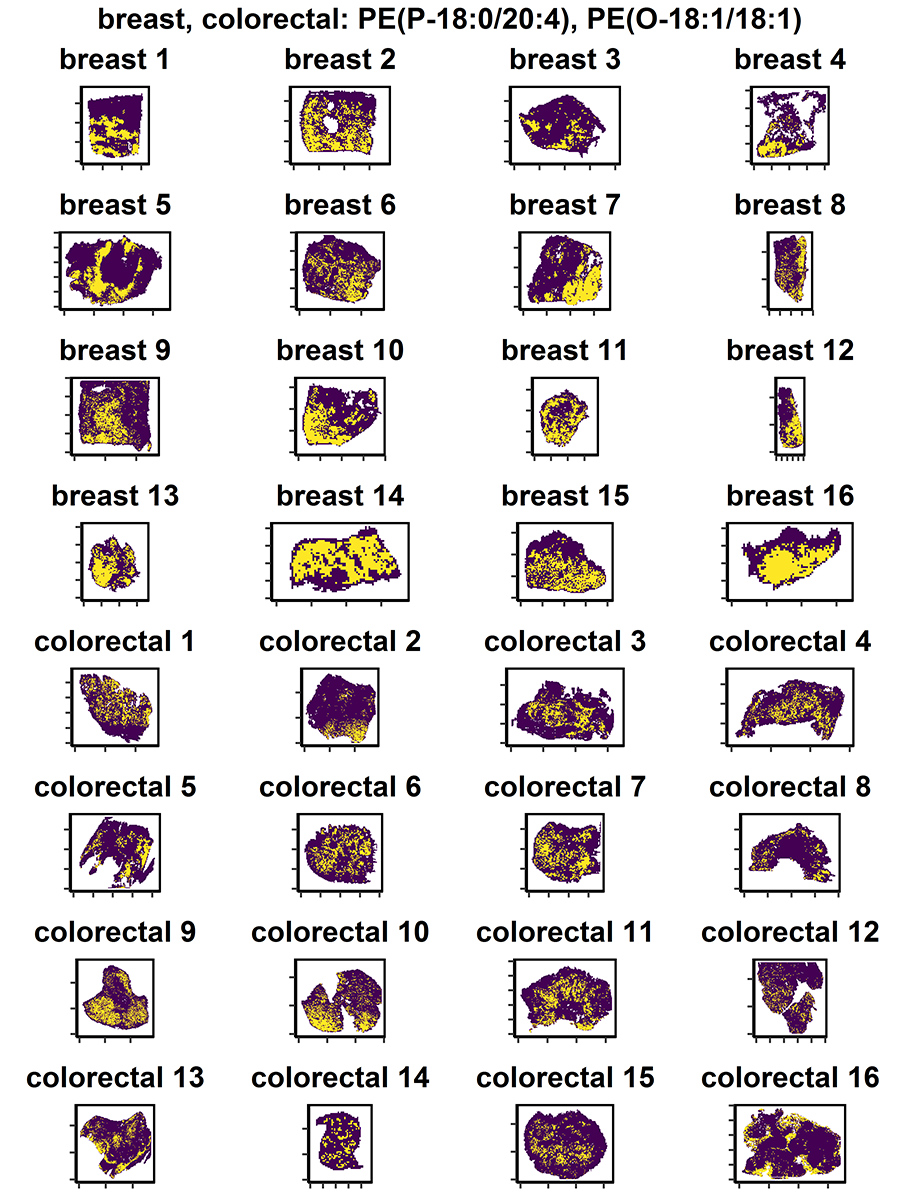

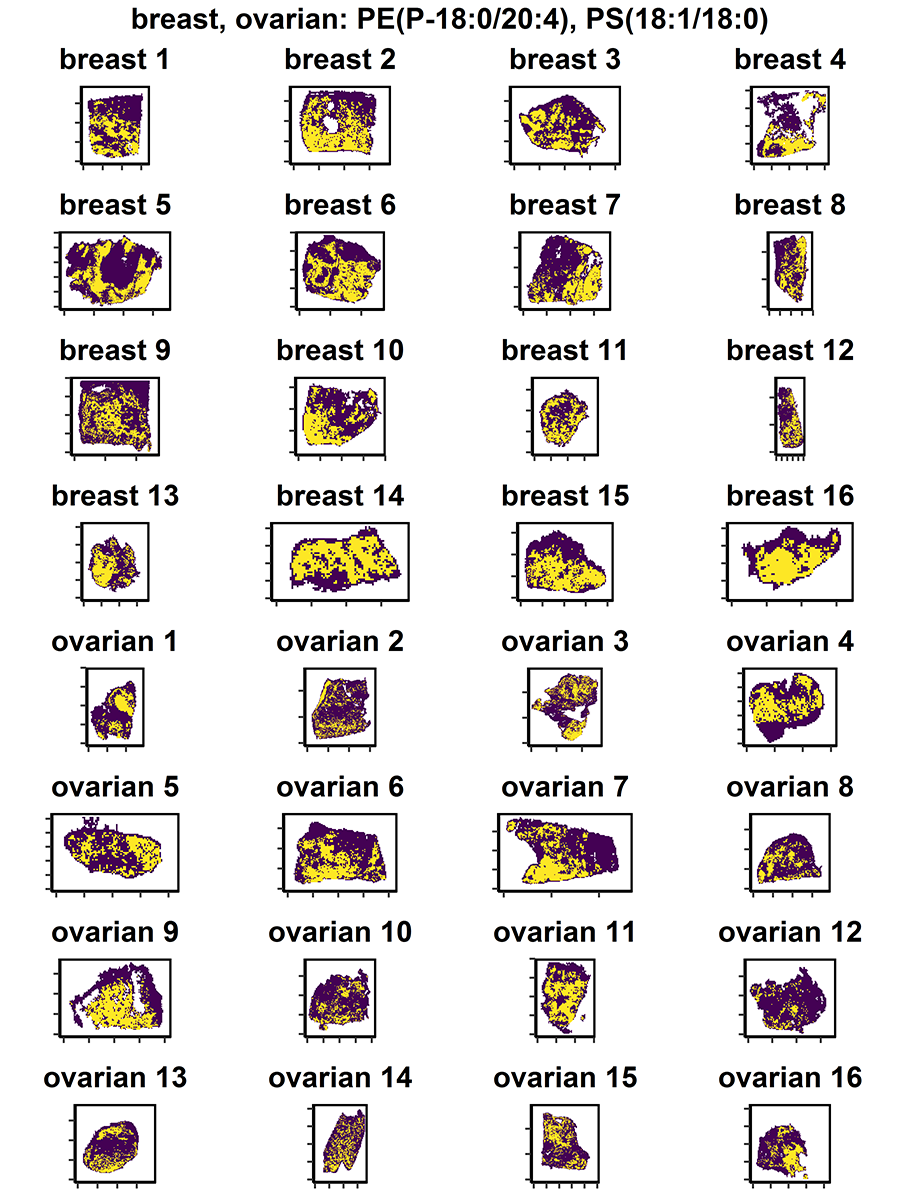

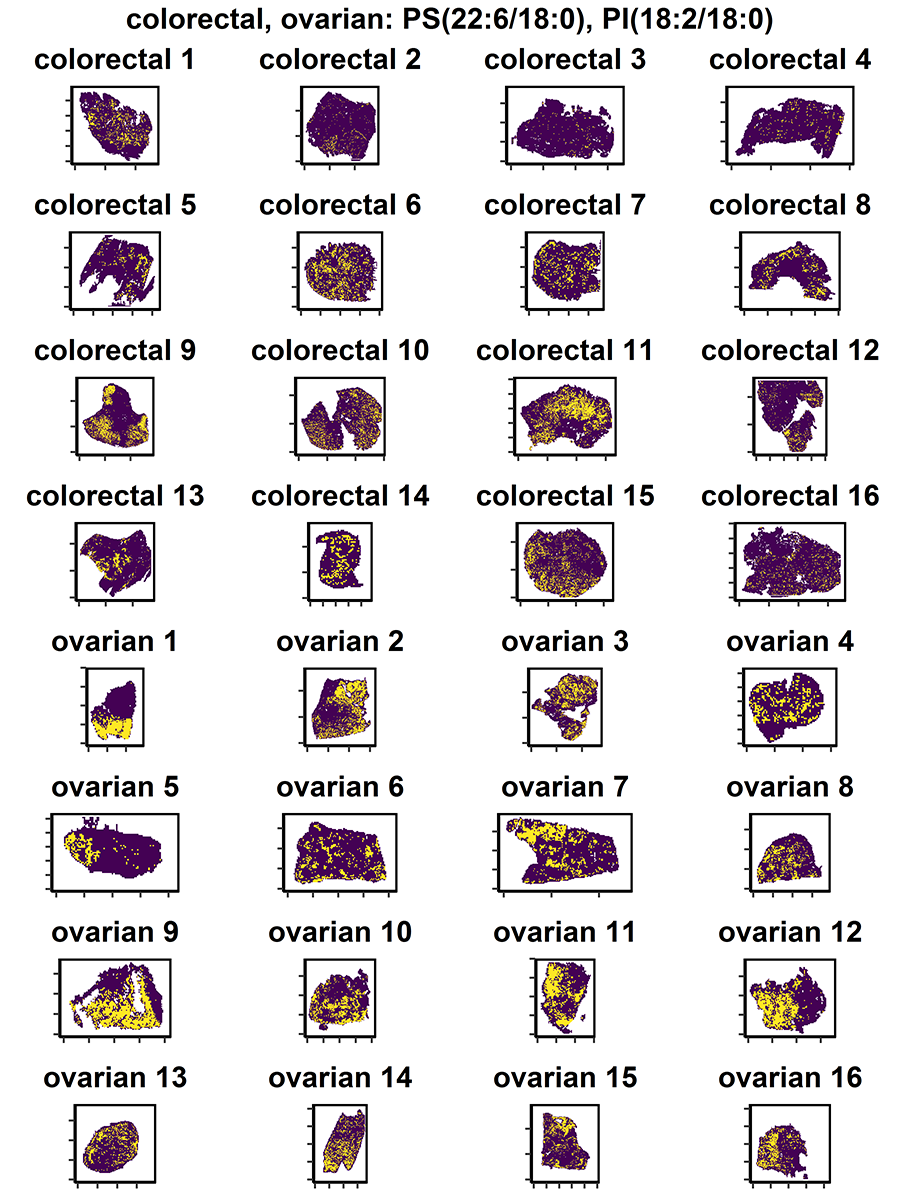

Figure S 6 – Clustering of the relative abundances of the ion pairs associated with the most significantly different correlations in the three contrasts. The yellow pixels represent the areas where the relative abundance of the two ions were higher than their average value.

Table S 1 – List of DESI-MS parameters used for the acquisition of the imaging data of the three sets of tissue sections. The columns represent the tissue section thickness, the lateral spatial resolution of the imaging data, the solvent flow rate and the nitrogen gas pressure used for running DESI-MS, respectively.

| **ID** | **Thickness (µm)** | **Resolution (µm)** | **Flow rate (µl/min)** | **Pressure (bar)** |
| --- | --- | --- | --- | --- |
| **Breast Samples** | | | | |
| 0502202A | 15 | 100 | 1.5 | 7 |
| 502263 | 15 | 100 | 1.5 | 7 |
| 602115 | 15 | 100 | 1.5 | 7 |
| 602167 | 15 | 100 | 1.5 | 7 |
| 702006 | 15 | 100 | 1.5 | 7 |
| 702233A | 15 | 100 | 1.5 | 7 |
| 702273 | 15 | 100 | 1.5 | 7 |
| 802086A | 15 | 100 | 1.5 | 7 |
| 802157A | 15 | 100 | 1.5 | 7 |
| 802183 | 15 | 100 | 1.5 | 7 |
| 802186 | 15 | 100 | 1.5 | 7 |
| 0902103A | 15 | 100 | 1.5 | 7 |
| 1102011A | 15 | 100 | 1.5 | 7 |
| 1102052A | 15 | 100 | 1.5 | 7 |
| 1102116 | 15 | 100 | 1.5 | 7 |
| 1102138 | 15 | 100 | 1.5 | 7 |
| BRB01S | 15 | 100 | 1.5 | 7 |
| BRB03N | 15 | 90 | 1.5 | 7 |
| BRB04N | 15 | 65 | 1.5 | 7 |
| BRB06E | 15 | 100 | 1.5 | 7 |
| BRB12S | 15 | 100 | 1.5 | 7 |
| BRB13D | 15 | 100 | 1.5 | 7 |
| CX12035 | 15 | 100 | 1.5 | 7 |
| CX12068 | 15 | 100 | 1.5 | 7 |
| CX12230 | 15 | 100 | 1.5 | 7 |
| CXH12000200 | 15 | 100 | 1.5 | 7 |
| CXH12000251 | 15 | 100 | 1.5 | 7 |
| **Colorectal Samples** | | | | |
| A15 | 10 | 100 | 1.5 | 7 |
| A20 | 10 | 110 | 1.5 | 7 |
| A22 | 10 | 100 | 1.5 | 7 |
| A26 | 10 | 100 | 1.5 | 7 |
| A28 | 10 | 100 | 1.5 | 7 |
| A33 | 10 | 100 | 1.5 | 7 |
| A37 | 10 | 85 | 2.5 | 7 |
| A49 | 10 | 115 | 1.5 | 7 |
| A52 | 10 | 80 | 1.5 | 7 |
| A53 | 10 | 90 | 1.5 | 5 |
| A56 | 10 | 125 | 1.5 | 5 |
| A57 | 10 | 90 | 1.5 | 5 |
| A62 | 10 | 100 | 1.5 | 7 |
| A64 | 10 | 85 | 1.5 | 5 |
| A74 | 10 | 100 | 1.5 | 7 |
| **Ovarian Samples** | | | | |
| *All* | 10 | 100 | 1.5 | 7 |

Table S 2 – List of parameters used for MALDIquant peak matching, within the MS image and between MS images, respectively.

|  | **Method** | **Tolerance** | **Min. Freq. Threshold** |
| --- | --- | --- | --- |
| **Within MS image** | strict | 10 ppm | 0.5% |
| **Between MS images** | strict | 20 ppm | 100% |

Table S 3 – List of SPUTNIK parameters used for filtering the tissue-unrelated peaks.

| **Global similarity filter** | |
| --- | --- |
| **Reference** | Binary ROI (k-means2) |
| **Method** | Pearson’s correlation |
| **Threshold** | 0 |
| **Pixel count filter** | |
| **Reference** | Binary ROI (k-means2) |
| **Min. number of connected pixels** | 9 |
| **Aggressiveness** | 1 |

Table S 4 – Annotations of the common m/z values detected after peak matching. Annotation based on MS/MS are reported in the last column: YES, MS/MS was used to identify the molecule, YES (N/A), the molecule could not be identified by its MS/MS spectrum; NO, the molecule was identified using only the m/z value.

| **Obs. *m/z*** | **Theor. *m/z*** | **Error (ppm)** | **Sum formula** | **Adduct** | **Compound** | **MS/MS** |
| --- | --- | --- | --- | --- | --- | --- |
| 259.2422 | 259.2431 | 3.4 | C19H32 | M-H- | Nonadecatetraene | YES (N/A) |
| 279.2321 | 279.233 | 3.3 | C18H32O2 | M-H- | Linoleic acid (C18:2) | YES (N/A) |
| 303.2321 | 303.233 | 2.9 | C20H32O2 | M-H- | Eicosatetraenoic acid (C20:4) | YES (N/A) |
| 305.2478 | 305.2486 | 2.6 | C20H34O2 | M-H- | Eicosatrienoic acid (C20:3) | YES (N/A) |
| 307.2635 | 307.2643 | 2.7 | C20H36O2 | M-H- | Eicosadienoic acid (C20:2) | YES (N/A) |
| 309.2791 | 309.2799 | 2.5 | C20H38O2 | M-H- | Eicosenoic acid (C20:1) | YES (N/A) |
| 327.2322 | 327.233 | 2.5 | C22H32O2 | M-H- | Docosatriynoic acid (C22:3) | NO |
| 331.2635 | 331.2643 | 2.4 | C22H36O2 | M-H- | Docosatetraenoic acid (C22:4) | YES (N/A) |
| 365.3417 | 365.3425 | 2.3 | C24H46O2 | M-H- | Nervonic acid (C24:1) | YES (N/A) |
| 419.2562 | 419.2568 | 1.4 | C21H41O6P | M-H- | Cyclic Phosphatidic acid (C18:0) | YES |
| 437.2666 | 437.2674 | 1.7 | C21H43O7P | M-H- | LysoPA(18:0) | YES |
| 464.3139 | 464.3146 | 1.6 | C23H48NO6P | M-H- | PE(P-18:0) | YES (N/A) |
| 480.3089 | 480.3096 | 1.4 | C23H48NO7P | M-H- | LysoPE(18:0) | YES |
| 536.5043 | 536.5048 | 0.9 | C34H67NO3 | M-H- | N-Palmitoylsphingosine / Cer(d18:1/16:0) | YES |
| 646.6142 | 646.6144 | 0.3 | C42H81NO3 | M-H- | Ceramide (d18:1/24:1) | YES |
| 673.4812 | 673.4814 | 0.4 | C37H71O8P | M-H- | PA(18:1/14:0) / PA(16:0/16:1) | YES |
| 687.5444 | 687.5446 | 0.2 | C38H77N2O6P | M-H- | SM(d33:1) / PE-Cer(d36:1) | YES (N/A) |
| 698.513 | 698.513 | 0 | C39H74NO7P | M-H- | PE(P-16:0/18:2) | YES |
| 699.497 | 699.497 | 0 | C39H73O8P | M-H- | PA(18:1/18:1) / PA(18:0/18:2) | YES |
| 700.5286 | 700.5287 | 0.2 | C39H76NO7P | M-H- | PE(P-14:0/18:1) | YES |
| 701.5126 | 701.5127 | 0.1 | C39H75O8P | M-H- | PA(36:1) | NO |
| 714.5079 | 714.5079 | 0 | C39H74NO8P | M-H- | PE(14:0/18:2) | YES |
| 716.5236 | 716.5236 | 0 | C39H76NO8P | M-H- | PE(14:0/18:1) | YES |
| 718.5392 | 718.5392 | 0 | C39H78NO8P | M-H- | PE(16:0/16:0) | YES |
| 722.5131 | 722.513 | -0.1 | C41H74NO7P | M-H- | PE(P-16:0/20:4) | YES |
| 723.4968 | 723.497 | 0.3 | C41H73O8P | M-H- | PA(18:0/20:4) / PA(18:2/18:2) | YES |
| 725.5128 | 725.5127 | -0.1 | C41H75O8P | M-H- | PA(18:0/20:3) | YES |
| 726.5445 | 726.5443 | -0.3 | C41H78NO7P | M-H- | PE(O-18:2/18:1) | YES |
| 728.5601 | 728.56 | -0.1 | C41H80NO7P | M-H- | PE(O-18:1/18:1) | YES |
| 738.5082 | 738.5079 | -0.4 | C41H74NO8P | M-H- | PE(16:0/20:4) | YES |
| 740.5240 | 740.5236 | -0.5 | C41H76NO8P | M-H- | PE(18:1/18:2) | YES |
| 742.5394 | 742.5392 | -0.3 | C41H78NO8P | M-H- | PE(18:1/18:1) | YES |
| 744.5551 | 744.5549 | -0.3 | C41H80NO8P | M-H- | PE(18:0/18:1) | YES |
| 746.5139 | 746.513 | -1.2 | C43H74NO7P | M-H- | PE(P-16:0/22:6) | YES |
| 748.5300 | 748.5287 | -1.8 | C43H76NO7P | M-H- | PE(P-18:1/20:4) | YES |
| 750.5443 | 750.5443 | 0.1 | C43H78NO7P | M-H- | PE(P-18:0/20:4) | YES |
| 752.5603 | 752.56 | -0.4 | C43H80NO7P | M-H- | PE(P-18:0/20:3) | YES |
| 762.5083 | 762.5079 | -0.5 | C43H74NO8P | M-H- | PE(16:0/22:6) | YES |
| 764.5237 | 764.5236 | -0.1 | C43H76NO8P | M-H- | PE(18:1/20:4) | YES |
| 766.5393 | 766.5392 | -0.1 | C43H78NO8P | M-H- | PE(18:0/20:4) | YES |
| 768.5554 | 768.5549 | -0.7 | C43H80NO8P | M-H- | PE(18:0/20:3) | YES |
| 769.5026 | 769.5025 | -0.2 | C42H75O10P | M-H- | PG(18:2/18:2) | YES |
| 770.5708 | 770.5705 | -0.4 | C43H82NO8P | M-H- | PE(18:1/20:1) / PE(18:0/20:1) | YES |
| 771.5183 | 771.5182 | -0.2 | C42H77O10P | M-H- | PG(18:1/18:2) | YES |
| 772.5867 | 772.5862 | -0.7 | C43H84NO8P | M-H- | PE(38:1) | NO |
| 773.5337 | 773.5338 | 0.2 | C42H79O10P | M-H- | PG(18:1/18:1) | YES |
| 774.5472 | 774.5443 | -3.8 | C45H78NO7P | M-H- | PE(P-18:0/22:6) | YES |
| 776.5619 | 776.56 | -2.4 | C45H80NO7P | M-H- | PE(P-18:0/22:5) | YES |
| 778.5761 | 778.5756 | -0.6 | C45H82NO7P | M-H- | PE(P-18:0/22:4) / PE(P-20:0/20:4) | YES |
| 786.5291 | 786.5291 | 0 | C42H78NO10P | M-H- | PS(18:2/18:0) | YES |
| 788.5448 | 788.5447 | -0.2 | C42H80NO10P | M-H- | PS(18:1/18:0) | YES |
| 792.5553 | 792.5549 | -0.5 | C45H80NO8P | M-H- | PE(18:0/22:5) | YES |
| 794.5709 | 794.5705 | -0.6 | C45H82NO8P | M-H- | PE(18:0/22:4) | YES |
| 810.5291 | 810.5291 | 0 | C44H78NO10P | M-H- | PS(20:4:18:0) | YES |
| 812.5454 | 812.5447 | -0.8 | C44H80NO10P | M-H- | PS(20:3/18:0) | YES |
| 816.5768 | 816.576 | -1 | C44H84NO10P | M-H- | PS(18:1/20:0) | YES |
| 820.5626 | 820.5629 | 0.4 | C44H84NO8P | M+Cl- | PC(18:1/18:1) | YES |
| 833.5188 | 833.5186 | -0.2 | C43H79O13P | M-H- | PI(18:2/16:0) | YES |
| 834.5308 | 834.5291 | -2 | C46H78NO10P | M-H- | PS(22:6/18:0) | YES |
| 857.5186 | 857.5186 | 0 | C45H79O13P | M-H- | PI(20:4/16:0) | YES |
| 859.5352 | 859.5342 | -1.2 | C45H81O13P | M-H- | PI(36:3) | NO |
| 861.5501 | 861.5499 | -0.3 | C45H83O13P | M-H- | PI(18:2/18:0) | YES |
| 883.5344 | 883.5342 | -0.2 | C47H81O13P | M-H- | PI(20:4/18:1) | YES |
| 885.5501 | 885.5499 | -0.2 | C47H83O13P | M-H- | PI(20:4/18:0) | YES |

Table S 5 – List of the five largest correlated pairs of ions in the three tissue types.

| **Ion 1** | **Ion 2** | **Correlation** |
| --- | --- | --- |
| **Breast** | | |
| PA(36:1) | PS(18:1/18:0) | 0.91964294 |
| PG(18:1/18:2) | PG(18:1/18:1) | 0.85387172 |
| PE(P-18:0/20:4) | PE(P-18:1/20:4) | 0.82841959 |
| PE(P-18:1/20:4) | PE(P-18:0/20:4) | 0.82781521 |
| PE(P-16:0/20:4) | PE(P-18:0/20:4) | 0.81827409 |
| **Colorectal** | | |
| PA(36:1) | PS(18:1/18:0) | 0.89393631 |
| PE(O-18:2/18:1) | PE(O-18:1/18:1) | 0.84356314 |
| PI(18:2/16:0) | PI(18:2/18:0) | 0.83477204 |
| PE(18:1/18:1) | PE(18:0/18:1) | 0.79990005 |
| PE(18:1/18:2) | PE(18:1/18:1) | 0.79349843 |
| **Ovarian** | | |
| PA(36:1) | PS(18:1/18:0) | 0.87004032 |
| PG(18:1/18:2) | PG(18:1/18:1) | 0.7705973 |
| PE(18:1/18:1) | PE(18:0/18:1) | 0.75112221 |
| PE(18:0/18:1) | PE(18:1/20:1) / PE(18:0/20:1) | 0.73529541 |
| PE(18:1/20:1) / PE(18:0/20:1) | PC(18:1/18:1) | 0.71865099 |

Table S 6 –Statistically significant different ion pair correlations obtained for the three contrasts are reported with the sum formulas corresponding to the m/z difference between the selected ions.

| **Molecule 1** | **Molecule 2** | **Sum formula** | **m/z 1** | **m/z 2** | **Δ m/z** | **Δ Sum formula** |
| --- | --- | --- | --- | --- | --- | --- |
| **Breast - Colorectal** | | | | | | |
| PE(P-16:0/20:4) | PE(O-18:1/18:1) | C41H74NO7P , C41H80NO7P | 722.513 | 728.56 | 6.047 | (+H6 (3 double bonds)) |
| PE(P-14:0/18:1) | PE(P-18:0/20:4) | C39H76NO7P, C43H78NO7P | 700.5287 | 750.5443 | 50.0156 | (+C4H2) |
| PE(18:0/18:1) | PE(18:0/22:5) | C41H80NO8P, C45H80NO8P | 744.5549 | 792.5549 | 48 | (+C4) |
| PE(18:0/22:4) | PC(18:1/18:1) | C45H82NO8P, C44H84NO8P | 794.5705 | 820.5629 | 25.9924 | (+CH2) |
| PS(18:1/20:0) | PS(22:6/18:0) | C44H84NO10P, C46H78NO10P | 816.576 | 834.5291 | 17.9531 | (+C2 –H6) |
| C20:4 | PE(O-18:1/18:1) | C20H32O2, C41H80NO7P | 303.233 | 728.56 | 425.327 | (+C21H48NO5P) |
| PE(O-18:1/18:1) | PS(22:6/18:0) | C41H80NO7P, C46H78NO10P | 728.56 | 834.5291 | 105.9691 | (+C5O3 –H2) |
| PE(18:0/18:1) | PE(18:0/22:4) | C41H80NO8P, C45H82NO8P | 744.5549 | 794.5705 | 50.0156 | (+C4H2) |
| PE(18:0/18:1) | PE(18:0/20:4) | C41H80NO8P, C43H78NO8P | 744.5549 | 766.5392 | 21.9843 | (+C2 –H2) |
| PE(O-18:1/18:1) | PE(P-18:1/20:4) | C41H80NO7P,  C43H76NO7P | 728.56 | 748.5287 | 19.9687 | (+C2 –H2) |
| **Breast - Ovarian** | | | | | | |
| PE(P-16:0/20:4) | PS(18:1/18:0) | C41H74NO7P, C42H80NO10P | 722.513 | 788.5447 | 66.0317 | (+CH6O3) |
| PA(36:1) | PE(P-18:0/20:4) | C39H75O8P, C41H74NO7P | 701.5127 | 722.513 | 21.0003 | (+C2 -OH) |
| PE(P-18:0/20:4) | PE(P-18:0/20:3) | C41H74NO7P, C43H80NO7P | 722.513 | 752.56 | 30.047 | (+C2H6) |
| PE(P-14:0/18:1) | PE(P-18:1/20:4) | C39H76NO7P,  C43H76NO7P | 700.5287 | 748.5287 | 48 | (+C4) |
| PE(P-18:0/20:4) | PE(P-18:1/20:4) | C41H74NO7P, C43H76NO7P | 722.513 | 748.5287 | 26.0157 | (+C2H2) |
| PS(18:1/18:0) | PI(20:4/18:0) | C42H80NO10P, C47H83O13P | 788.5447 | 885.5499 | 97.0052 | (+C5H3NO3) |
| PE(P-18:1/20:4) | PI(20:4/16:0) | C43H76NO7P, C45H79O13P | 748.5287 | 857.5186 | 108.9899 | (+C2H3NO6) |
| PE(P-16:0/22:6) | PS(18:1/18:0) | C43H74NO7P, C42H80NO10P | 746.513 | 788.5447 | 42.0317 | (-C +H6O3) |
| PA(36:1) | PI(20:4/18:0) | C39H75O8P, C47H83O13P | 701.5127 | 885.5499 | 184.0372 | (+C8H8O5) |
| PA(36:1) | PE(P-18:0/20:4) | C39H75O8P, C43H78NO7P | 701.5127 | 750.5443 | 49.0316 | (+C4H2 -O ) |
| **Colorectal – Ovarian** | | | | | | |
| PS(22:6/18:0) | PI(18:2/18:0) | C46H78NO10P, C45H83O13P | 834.5291 | 861.5499 | 27.0208 | (-CN /+H5O3) |
| C22:4 | PE(P-18:0) | C22H36O2, C23H48NO7P | 331.2643 | 464.3146 | 133.0503 | (+CH12O5P) |
| PS(20:3/18:0) | PI(18:2/18:0) | C44H80NO10P, C45H83O13P | 812.5447 | 861.5499 | 49.0052 | (+CH3O3 /-N) |
| Cer(d18:1/16:1) | PE(18:0/22:4) | C34H67NO3, C45H82NO8P | 536.5048 | 794.5705 | 258.0657 | (+C11H15O5P) |
| PS(20:3/18:0) | PI(18:2/16:0) | C44H80NO10P, C43H79O13P | 812.5447 | 833.5186 | 20.9739 | (+NO3/ -CH) |
| PE(P-18:0) | PS(22:6/18:0) | C23H48NO6P, C46H78NO10P | 464.3146 | 834.5291 | 370.2145 | (+C23H30O4) |
| PE(16:0/16:0) | PI(20:4/18:0) | C39H78NO8P,  C47H83O13P | 718.5392 | 885.5499 | 167.0107 | (+C8H5O5 / -N) |
| C22:4 | PI(18:2/18:0) | C22H36O2, C45H83O13P | 331.2643 | 861.5499 | 530.2856 | (+C23H47O11P) |
| PE(18:0/20:4) | PI(18:2/16:0) | C43H78NO8P,  C43H79O13P | 766.5392 | 833.5186 | 66.9794 | (+HO5 /-N) |
| PE(P-18:0) | PA(18:0/20:4) /  PA(18:2/18:2) | C23H48NO6P, C41H73O8P | 464.3146 | 723.497 | 259.1824 | (+C18H25O2/ -N) |
